# Critical-Size Defect Tibialis Anterior (TA) Muscle Regeneration using Ex-Vivo Mice Hindlimbs Culturing under Dynamic Mechanical Loading

**DOI:** 10.1101/2024.10.16.618565

**Authors:** Diego Jacho, James Huynh, Emily Crowe, Agustin Rabino, Mine Yildirim, Piotr J. Czernik, Beata Lecka-Czernik, Rafael Garcia-Mata, Eda Yildirim-Ayan

## Abstract

In this study, we introduced an innovative computer-controlled ex vivo mice hindlimb culturing platform operating under dynamic loading, coupled with injectable cell-laden nanofibrous matrix (PNCOL), to investigate tissue response and therapeutic outcomes in critical size defect tibialis anterior (TA) muscle regeneration. The combination of mechanical stimulation and cell therapy offers a distinctive opportunity to delve into the regenerative rehabilitation field and create sustainable solutions in musculoskeletal (MSK) tissue regeneration. The application of mechanical loading on the whole mice hindlimbs increased total bone area and marrow area suggesting an increase in periosteal bone formation and resorption on the endosteal surface. Viability assessments confirmed the sustained culturing of the samples throughout the study. Then, the effect of mechanical loading and PNCOL injection on muscle regeneration at the TA defect site was evaluated. Histological analyses revealed enhanced muscle regeneration in PNCOL-treated hindlimbs. Structural analysis of the defect area through scanning electron microscopy (SEM) showed regeneration of ECM fibers at the defect site in PNCOL-treated groups. An analysis of cytokine levels in conditioned media at the end experiment showed changes in the number of proteins with the role in wound healing, muscle regeneration WNT, and IGF-1 signaling suggesting an anabolic effect of mechanical stimulation on muscle and bone. Similarly, gene expression analysis showed a significant upregulation of PAX7, Mrf4, MYF5, and TGFβ1 mRNA levels, indicating enhanced muscle regeneration after coupled mechanical loading and PNCOL treatments. Lastly, immunostaining showed an increase in tissue regeneration and anti-inflammatory response (CD206) in PNCOL-treated groups. Overall, the ex vivo hindlimb organ culturing platform– maintained tissue functions under mechanical loading, while PNCOL treatment promoted muscle tissue regeneration and reduced inflammation. These findings demonstrated the potential of multidimensional approaches for enhancing therapeutic outcomes in MSK disorders. In addition, this study aligns with the growing emphasis on minimizing the number of animals used in research and developing a robust sense of responsible animal experimentation through introducing dynamic ex-vivo muscle organ culturing platform.

## 1. Introduction

Musculoskeletal (MSK) tissue disorders stand as the most prevalent physiological diseases worldwide, affecting approximately 1.71 billion people(1-4). The exploration of novel methodologies specifically through cell therapy and mechanical loading in forms of mechanotherapy(5) holds promise for enhancing MSK tissue regeneration. There has been an increase in autologous mesenchymal stem cell (MSC) utilization for various MSK injury treatments (6-8). To date, bone marrow-derived MSCs are the most frequently used cell source for MSK tissue repair by delivering them to the area of interest via a syringe or other minimally invasive techniques(9, 10). However, the injection of MSCs into the defect area comes with several challenges including post-injection cell death (9, 10) due to the harsh immune environment at the defect site and possible migration of the MSCs to the other areas subsequently creating ectopic tissue formations. To prevent these challenges, the MSCs are encapsulated within a synthetic and/or natural matrix which serves as a temporary home for cells until tissue regeneration and as a cell barrier matrix for harsh outside environments (10-12).

Utilizing synthetic and/or natural matrix along with the cell therapy is particularly important for critical - sized defects with a size ranging from 1 to 3 cm(13) and volumetric muscle loss (VML)(14). The natural tissue repair mechanism is overwhelmed in regenerating large tissue structures for critical-sized bone defects and VLM cases. To address this, cellular or acellular synthetic and/or natural matrices are designed as filler materials, not only for carrying and protecting the encapsulated cells, serving as an inductive template to recruit endogenous stem/progenitor cells to injury sites for repair(15-17) but also increasing mechanical stability at the defect site(8). For instance, Zhang Z., et al. (9). designed a conductive porous nanocomposite three-dimensional (3D) microgels loaded with myoblast which significantly improved the generation of new muscle fibers, blood vessels, and anti-inflammatory properties at a mouse VML injury site. Numerous seminal studies discuss the design criterion of synthetic and/or natural matrices synthesis process for bone and muscle regeneration(18-21); however, their performances in functional bone and muscle regenerations have yielded highly variable results (22-25) with limited in vivo application success.

The translation of in vitro success to in vivo applications is often hindered because many studies are conducted on two-dimensional (2D) surfaces (well plate and Petri dishes) without accounting for the surrounding extracellular matrix (ECM) and/or the dynamic mechanical environment of bones and muscles during culturing, which are crucial for functional muscle and bone regeneration and tissue recoveries(26). To address this gap, in vivo studies have been conducted to take advantage of the cell-matrix interaction within the 3D environment under physiological mechanical loading conditions. For instance, Kim J. et al. (27) explored tibialis anterior (TA) VML injuries in a rat model using allogenic decellularized skeletal muscle scaffolds, revealing the age, stiffness, and size of scaffolds as obstacles to regenerative medicine strategies targeting VML injury. Similarly, Sicari B. et al. (24) used an in vivo VML model to assess the efficacy of surgically placed inductive biologic scaffold material on mouse quadriceps, highlighting the therapeutic potential of a reproducible animal model to study VML. While in vivo studies have numerous superiorities over in vitro studies, they have their limitations as well including not being cost-efficient and being inherently inconsistent due to the biological variability and complexity of the animals (28, 29). In addition, variability in post-defect behavior among animals, where some may exhibit normal locomotion while others show limited movement or inactivity, can lead to inconsistent results, particularly in studies focusing on musculoskeletal tissues (30-32).

To circumvent these challenges, ex vivo MSK tissue culturing platforms with physiologically relevant mechanical stimulation have emerged. While in-vivo models still remain valuable for studying whole-organism physiology, ex-vivo platforms offer a powerful tool for dissecting the specific mechanisms that drive tissue regeneration. By isolating and controlling specific variables, ex-vivo methods enable researchers to obtain clearer insights into the underlying mechanisms of tissue response and regeneration. This reduces confounding variables and extraneous factors that may influence results in vivo, leading to a clearer understanding of the underlying mechanisms at play. The ex-vivo studies mitigate the complexities and ethical concerns associated with in vivo studies, aligning with the 4Rs principle (Reduction, Refinement, Replacement and Responsibility) (33). In sum compared to in-vivo studies, the ex-vivo models can replace certain in-vivo studies, reduce the number of animals required, and refine the methods to minimize animal suffering and improve animal welfare, and help to deliver faster, more reproducible, cost-effective and responsible results (34-36).

In addition, ex-vivo models, unlike 2D studies, preserve crucial 3D cell-cell and cell-matrix interactions, enabling the examination of native cells and tissues within their natural physiological environment under physiological mechanical loading with a fraction of the cost of in vivo studies and with higher consistency (37-41). For instance, Passipieri J. et al. (6, 42) developed an implantable tissue-engineered construct that facilitated substantial recovery in a VML injury model using ex vivo muscle extracted from rodents. Similarly, Sargent M. et al(43) used muscle tissue from Sprague Dawley rats to study the response to physical stress, highlighting the efficacy of ex vivo models under mechanical stimulation in understanding the severity of muscle damage in MSK injuries. These ex-vivo studies and many others shed light on how bone and muscle regenerate under mechanical loading using various interventions (cell therapy, biomaterial-based therapies, etc.,); however, they isolate either bone or muscle based on their interest and only focus on one tissue during ex vivo culturing (43-46). Whole ex vivo organ models, encompassing both bone and muscle, can offer a unique advantage over single-tissue studies (39) by capturing the intricate interplay between bone and muscle physiology, thereby providing a more relevant understanding of MSK disorders.

Towards this end, in this study, a computer-controlled ex-vivo mice hindlimbs culturing platform operating under dynamic mechanical loading was developed to study the effect of cell-laden nanofibrous scaffold on a critical-size defect on tibialis anterior (TA) muscle regeneration. The ex vivo mice hindlimbs with defects at TA muscles were cultured for 7 days under 12% strain with 1Hz frequency. Following comprehensive histological, molecular, and proteomic analyses, it was proved that ex-vivo mice hindlimbs culturing platform maintained the physiological functions of the entire hindlimbs, and the TA defect treated with cell-laden nanofibrous scaffold, called PNCOL, demonstrated better regeneration than non-treated-defect-under-dynamic mechanical loading condition.

## 2. Results

### 2.1. Ex-Vivo Organ Culture Platform Efficiency on Whole Mice Hindlimbs

First, our examination of the whole mouse hindlimb within our organ culture provided significant insights into the effects of mechanical stimulation on bone structure. Utilizing micro-CT analysis, we focused on the bone area of the stimulated samples, particularly emphasizing mechanical loading stimulation. Our results, as presented in Figure 1, highlight changes in the tibia and femur midshaft cortical bone after stimulation. Following the stimulation, both the tibia and femur exhibited notable increases in total area of bone and marrow area compared to the control group (unstimulated). Total midshaft bone area increased in stimulated femur and tibia by 6% and 4%, respectively, while cortical thickness and bone area remained rather unchanged (±1%). However, the marrow area increased by 10% in both bones. These suggest increased bone formation on the periosteal surface, which is in the immediate contact with exercised muscle, and bone loss on the endosteal surface probably due to increased proinflammatory responses in the absence of blood supply and innervation. Overall, these results indicate that within the duration of the experiment, bone tissue was alive and responded to the biomechanical stimulation and altered environment.

**Figure 1.**
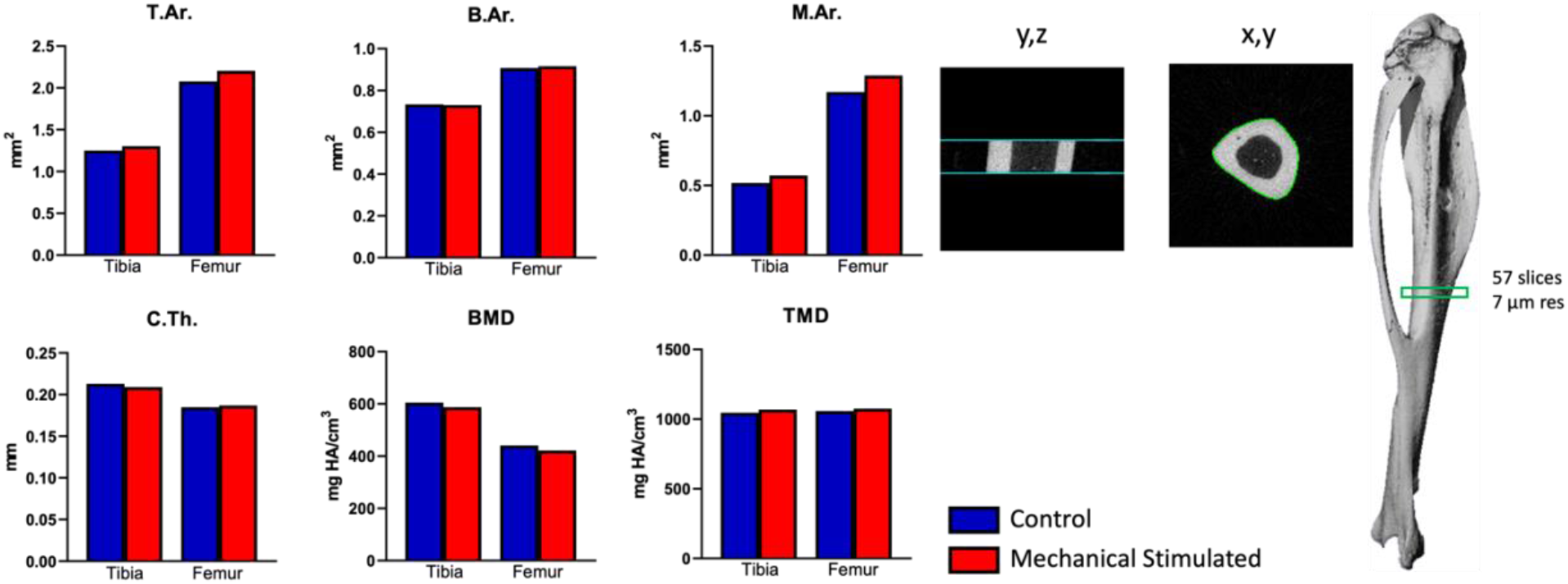
Micro-CT Analysis of Midshaft Femur at the End of Experiment. Measurements: Total area (bone plus marrow area) (T.Ar.), bone area (B.Ar.), marrow area (M.Ar.), mean cortical thickness (Ct. Th.), bone mineral density (BMD), and tissue mineral density (TMD).

### 2.2. Cell Viability within the Mice Hindlimb After Treatments

After demonstrating the efficiency of the organ culture platform in applying mechanical stimulation to the entire mice hindlimbs, 2-mm damage sites were created at the TA muscle and treated with myoblast-encapsulated polycaprolactone and collagen scaffolds, PNCOL (+). Briefly, PCL pellets were dissolved in a chloroform and methanol mixture, extruded through a needle, and collected as nanofibers, which were then homogenized, and oxygen-plasma treated. The treated nanofibers were mixed with a neutralized collagen type-I solution, and the mixture was used to encapsulate C2C12 mouse myoblast cells at a density of 10^6 cells/ml. All hindlimbs were exposed to the mechanical stimuli and damage sites without scaffolds, PNCOL (-), were used as control. After completing the mouse hindlimbs culture study, whole mouse hindlimbs were promptly removed from the culture setup to examine gross optical images between mechanically stimulated PNCOL (-) and PNCOL (+) hindlimbs.

Gross optical images revealed notable morphological contrasts between the hindlimbs treated with PNCOL and their untreated counterparts (Figure 2A). The PNCOL (+) group exhibited a fully restored, robust skeletal muscle structure with normal coloration. Conversely, in the control group, PNCOL (-) the puncture defect remained visible after 7 days of organ culturing. The cell viability was measured throughout the whole organ culture (7 days) period using the colorimetric-XTT assay. Figure 2B demonstrated that the cell viability was preserved throughout the organ culture period for both PNCOL (-) and PNCOL (+) samples. These results demonstrated that the whole organ culture was successfully kept viable in culture during the period of the study.

**Figure 2.**
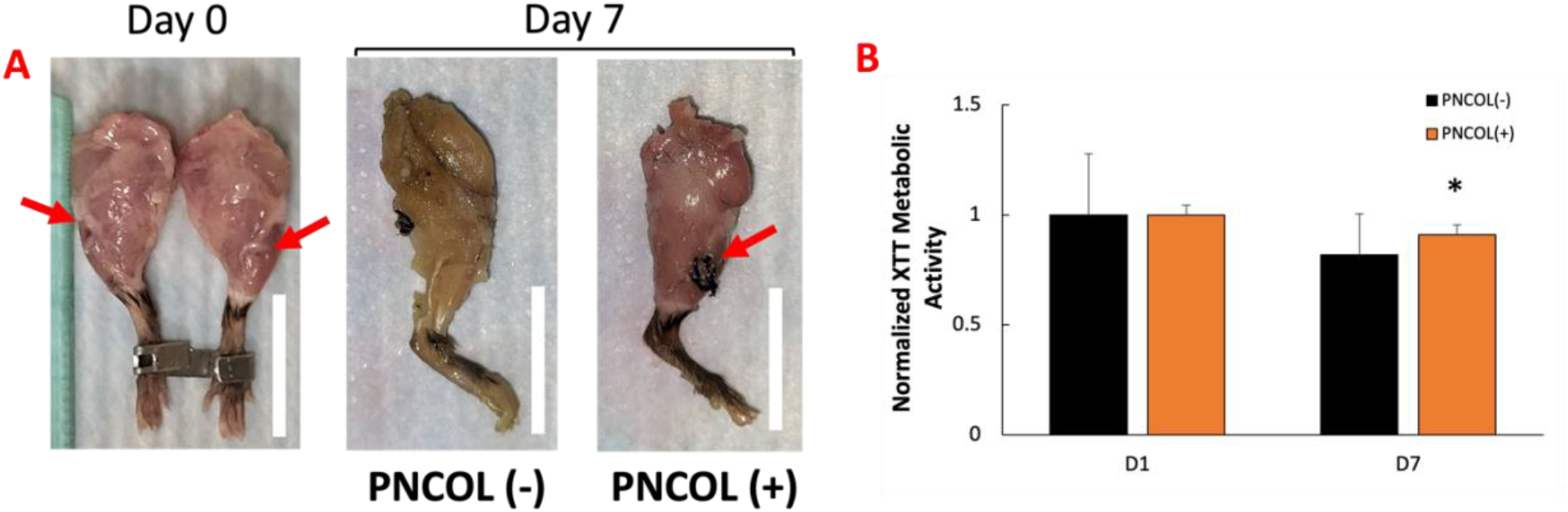
**A)** Representative optical Images of PNCOL (-) and PNCOL (+) mice hindlimbs after mechanical stimulation. Scale bar = 20mm, red arrow demonstrates damage site, and **B)** Cellular Metabolic Activity at days 1 and 7 after treatment and mechanical stimulation (n=4). (*) Indicates the statistical differences between control and experimental groups with p<0.05.

### 2.3. Structural and Morphological Stability of TA Muscle Defect Site After Treatments

To further explore the effects of PNCOL treatment and mechanical stimulation on TA muscle healing after damage, we conducted a histological analysis of the damage site. In the PNCOL (-) group, Figure 3 histology images revealed a notable absence of tissue regeneration and cell density. Furthermore, these images illustrated that the group receiving mechanical stimulation without treatment (PNCOL (-)) exhibited more space within the damaged area, resulting in no observable regeneration after 7 days of organ culturing. In contrast, the PNCOL (+) group displayed a significant increase in nuclei count in H&E staining, indicative of heightened satellite cell presence, essential for muscle regeneration (47, 48). Satellite cells, dormant muscle stem cells crucial for repair, proliferate and differentiate into mature muscle cells upon activation(49, 50). This elevated satellite cell presence in the PNCOL-treated group suggests superior regenerative potential compared to controls.

**Figure 3.**
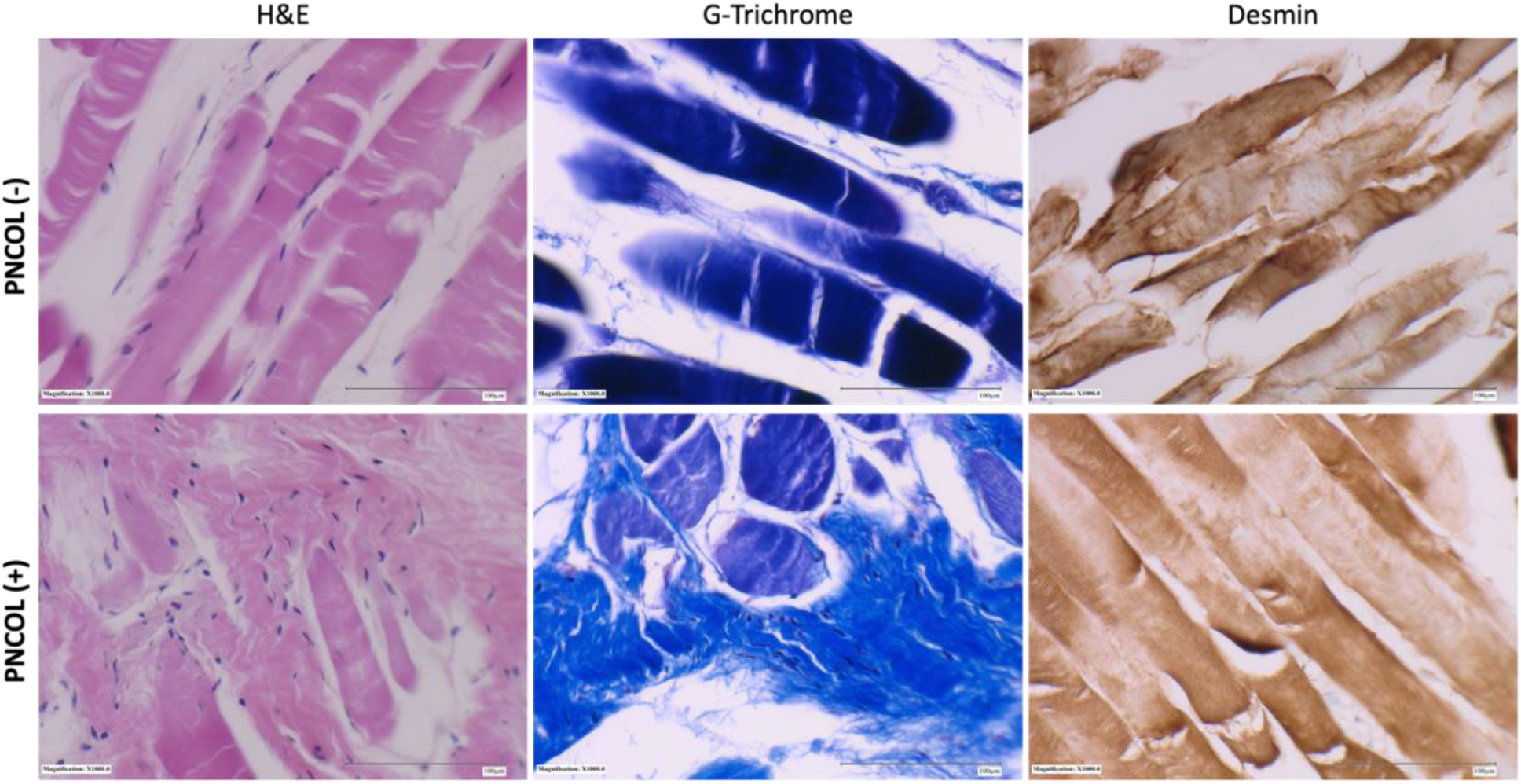
Representative histological images of the TA muscle stained with H&E, G-trichrome, and Desmin after 7 days of PNCOL treatment and mechanical stimulation (1000x). Scale bare = 100 µm (n=4).

Additionally, the slides were stained with desmin (DES), a muscle-specific protein crucial for maintaining muscle structure during myofibril regeneration (51, 52). Although quantitative measurements of diameter were not performed, qualitative observations of tissue morphology revealed notable differences between groups. The PNCOL (-) group muscle fibers displayed irregular shapes and a less cylindrical appearance compared to the more uniform and structured fibers observed in the PNCOL (+) treated samples. Additionally, the presence of increased space or ‘gaps’ within the damaged area in the PNCOL (-) group further indicated disrupted tissue organization and impaired regeneration. These muscle fibers were notably irregular and smaller in shape at the injury site than those in the PNCOL (+) group. Consequently, the PNCOL (-) group’s regenerating muscle fibers appeared smaller and less mature, indicating impaired muscle regeneration at the damage site in the PNCOL (-) group.

Injury recovery hinges on a balance of proteoglycan production; excessive levels can lead to scarring and tissue damage. Figure 4 illustrates heightened proteoglycan production in the PNCOL (+) group, indicative of robust muscle recovery and tissue regeneration, likely influenced by mechanical stimulation and PNCOL treatment (53-55). Conversely, the control group exhibited no significant regeneration in the defect area.

**Figure 4.**
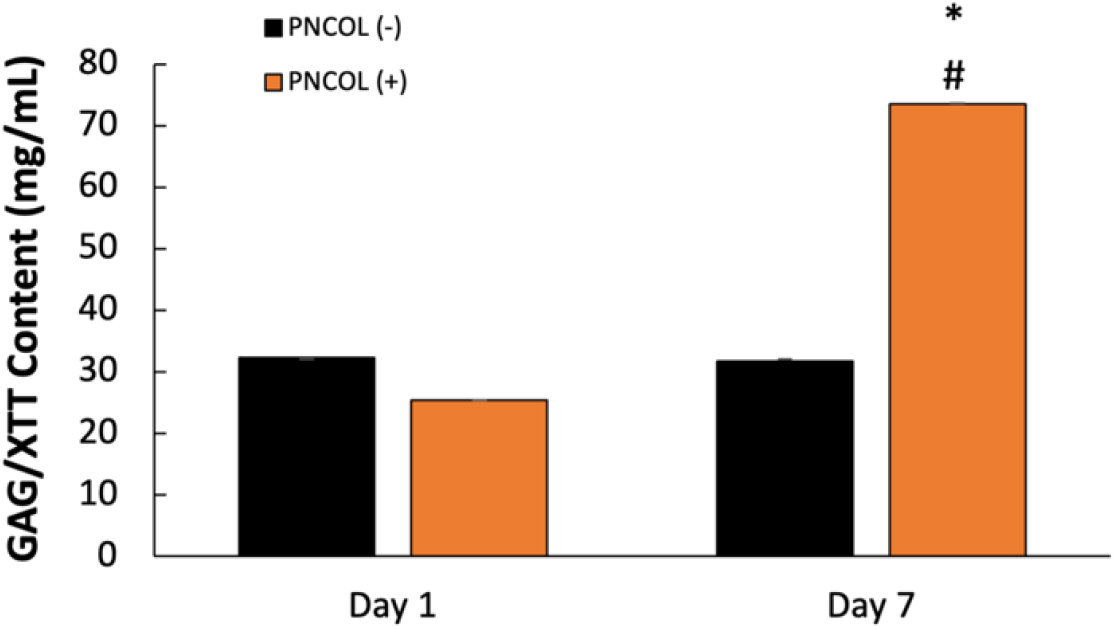
Proteoglycans production. GAG/XTT (mg/mL) content at days 1 and 7 after PNCOL treatment and mechanical stimulation(n=4). (*) Indicates the statistical differences between control and experimental groups with p<0.05, (#) indicates a statistical difference between experimental groups of the same day data point.

### 2.4. Matrix Organization and Muscle Fiber Alignment of TA Muscle Defect Site After Treatments

The representative SEM images of the TA defect treated with PNCOL and not treated are in Figure 5. The SEM images show the defect area (highlighted under white brackets) left untreated (PNCOL (-) group) shows a clear gap in the defect area, while The PNCOL (+) group demonstrates the ECM matrix production at the defect area. The PNCOL (+) SEM images showed more robust, compact fibers of muscle along the damage site while the PNCOL (-) group showed thin and wispy ECM projections between the interwoven meshes of collagen and tissue.

**Figure 5.**
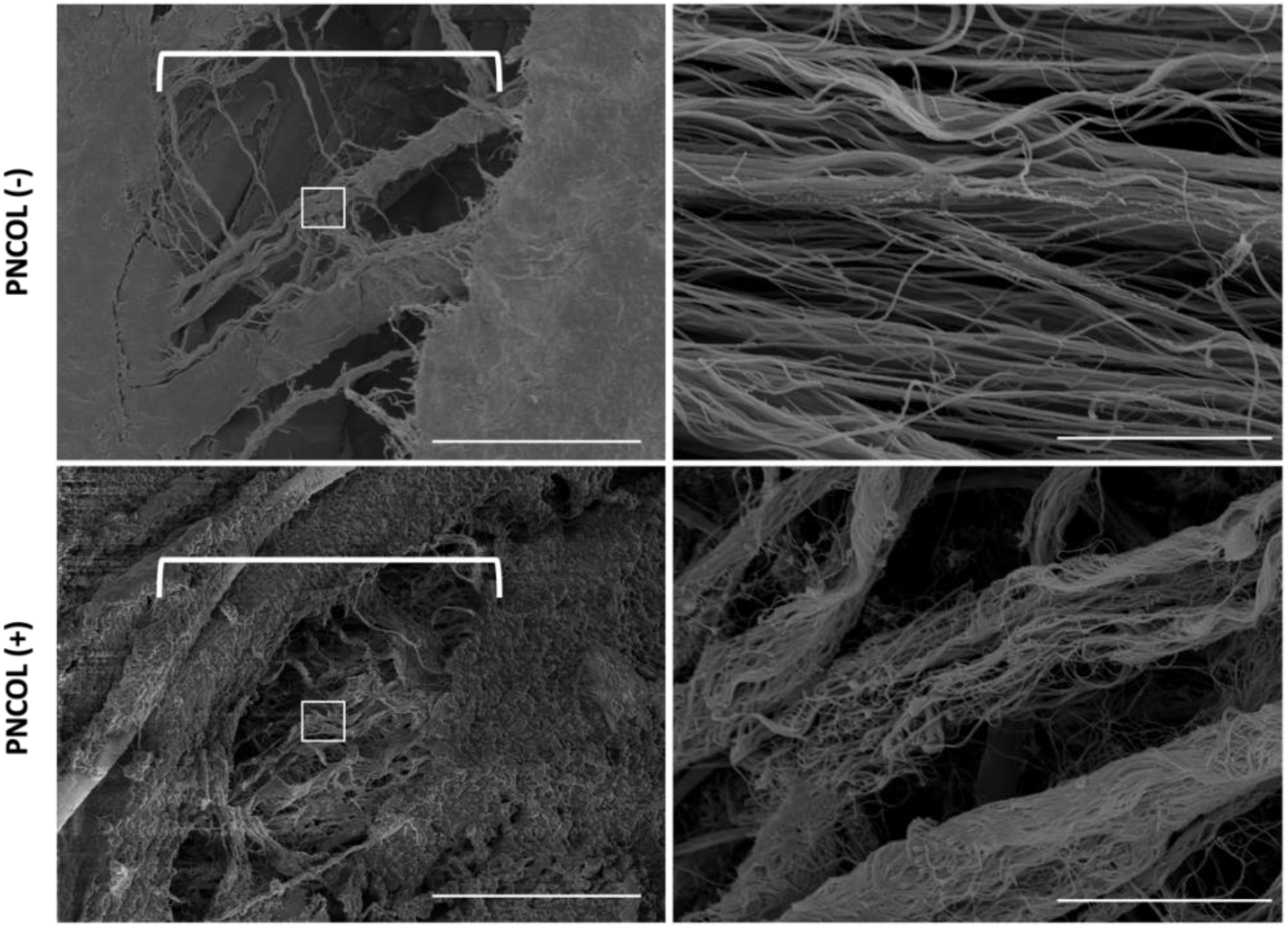
Representative scanning electron micrographs of PNCOL (-) and PNCOL (+) damage sites after 7 days of mechanical stimulation. The section under the brackets represents the damaged site area. Graphs on the right panel represent the zoom section of the damaged site. Scale bar is 50 µm for the left panel and 10 µm for the right panel. (n=4).

### 2.5. Cytokines and Chemokines Expression of Mice Hindlimbs Medium after Treatments

To investigate the effect of PNCOL and mechanical stimulation on the mice hindlimbs culture after treatments, we used proteome profiler arrays coated specifically with antibodies against mouse cytokines and chemokines. Media from both groups were collected and analyzed after 7 days of treatments. Several cytokines and chemokines were released to the media in response to both treatments (Figure 6). Among them factors involved in muscle regeneration (Adiponectin), inflammation and wound healing (CXCL1 and CXCL16), tissue remodeling (MMP2 and MMP3), and proteins that modulate IGF-1 signaling (IGFBP5 and IGFBP6). Consistent with bone response to both mechanical stimulation and PNCOL treatment, the levels of osteopontin and osteoprotegerin were increased. On the other hand, there was a decrease in the level of two inhibitors of the WNT signaling pathway (DKK1 and WISP1); the pathway which is anabolic for muscle and bone. Simultaneously, levels of several cytokines involved in inflammation, markers for chronic inflammation, and pro-inflammatory responses like ICAM-1, IFN-*γ*, and MMP9 were decreased in the PNCOL (+). Together the cytokine profile suggests a dynamic remodeling of the ECM, which is essential for tissue repair and regeneration, and is supported by the muscle and bone secretome. This aligns with the overall findings of our study, which demonstrates a modulation of immune responses and a favorable shift towards enhanced muscle regeneration in the PNCOL (+) group compared to the untreated counterparts. On the other hand, the cytokines and chemokines involved in cell regulation, proliferation, differentiation, and tissue remodeling were upregulated in the PNCOL (+) group compared to the PNCOL (-) group. Because detecting proteins secreted at a low level and diluted in the culture medium is difficult, we confirmed the major markers by RT-qPCR.

**Figure 6.**
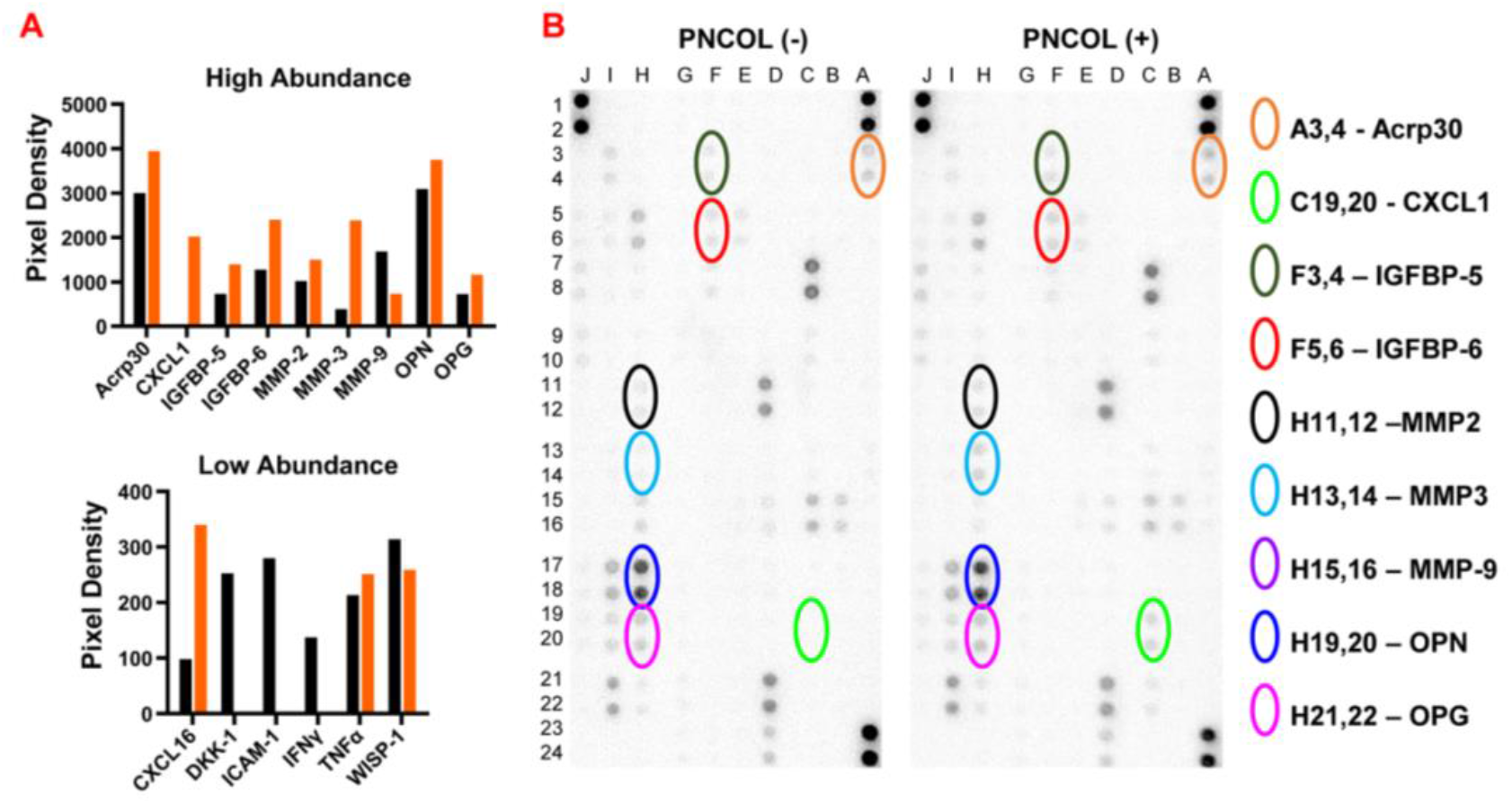
Cytokine levels in conditioned media (CM) collected from PNCOL (+) and PNCOL (-) samples on day 7. A) Differentially secreted high and low abundance cytokines in PNCOL (-) (black bars) and PNCOL (+) (orange bars) conditions. B) Membranes of the Proteome Cytokine Arrays incubated with CM from PNCOL (+) and PNCOL (-) samples with encircled spots corresponding to a high abundance of differentially expressed cytokines.

### 2.6. Gene Expression Analysis of TA Muscle Defect Site After Treatments

To understand the effect of mechanical stimulation and PNCOL treatment on muscle regeneration at the molecular level, a comprehensive gene expression analysis was conducted on PNCOL-treated and PNCOL (-) samples in all replicated experiments. Figure 7 shows the relative gene expression fold change of important muscle markers between PNCOL (-) and PNCOL (+) groups following 7 days of organ culturing. MRF4 and MYF5, genes encoding transcription factors crucial in regulating skeletal muscle development, were upregulated 3-fold and 5-fold, respectively in PNCOL (+) groups compared to PNCOL (-) groups. Also, PAX7 and *α*SMA, genes that play distinct but complementary roles in maintaining muscle repair and tissue remodeling, were upregulated in PNCOL (+) groups (Figure 7). The two important genes, myod, and myogenin, orchestrating the muscle cells’ differentiation into mature muscle fibers, only myogenin demonstrated an upregulation of almost 1.5-fold after the PNCOL treatment. Finally, the expression of VEGFA and TGFβ1 gene was upregulated almost 2-fold in the PNCOL (+) compared to PNCOL (-). The VEGFA and TGFβ1 roles are critical in promoting angiogenesis and regulating the inflammatory and fibrotic responses.

**Figure 7.**
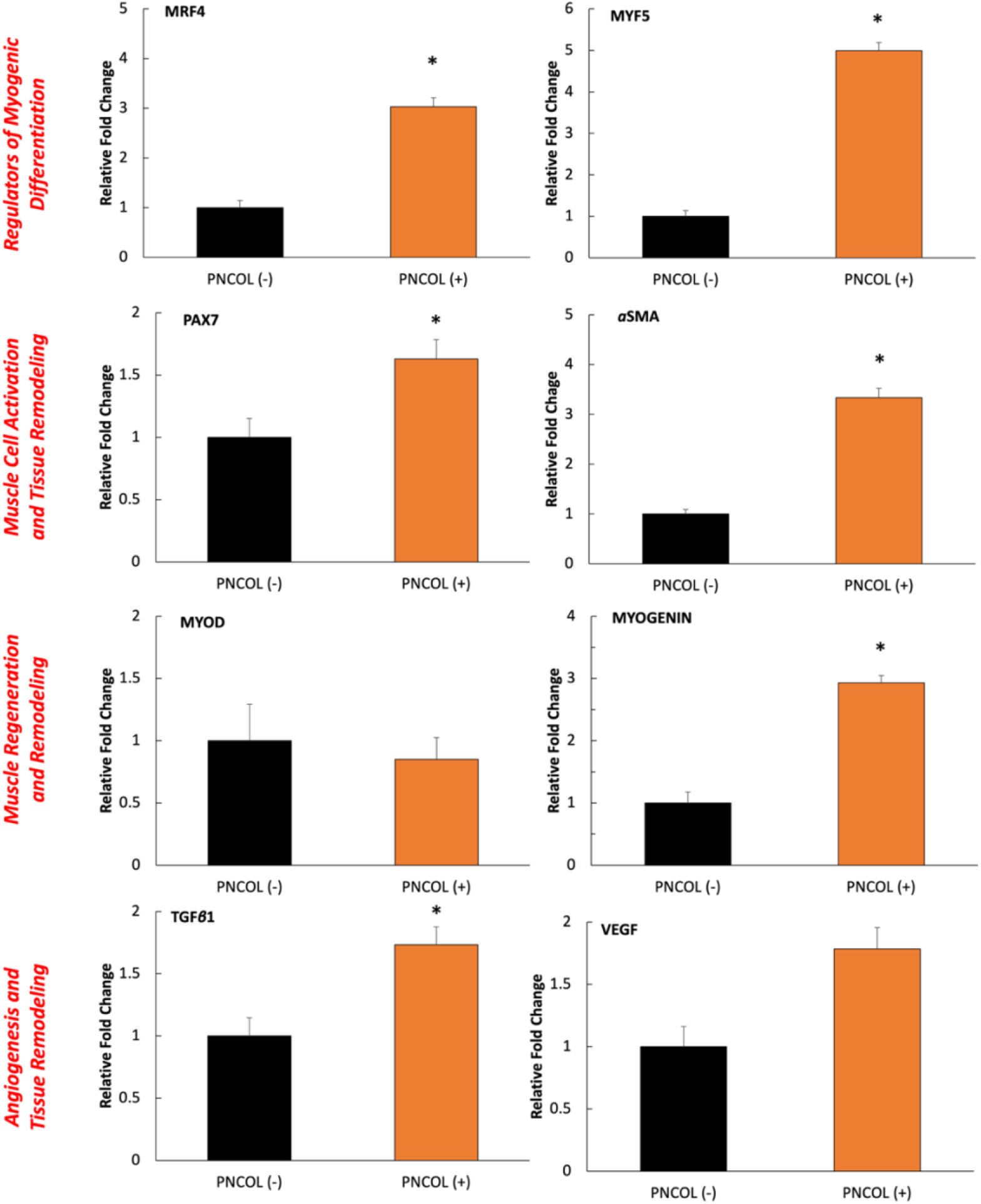
The relative fold changes of TA muscle markers of PNCOL (-), and PNCOL (+) samples, following 7 days of organ culture in a dynamic mechanical environment. All data were expressed as the mean and standard errors. *Indicates the statistical differences between groups with p<0.05, n=4; at least three technical replicates were used for gene expression analysis.

Next, genes involved in the inflammatory response were analyzed. The inflammatory response during muscle regeneration coordinates the clearing of damaged tissue, recruitment of immune cells, and initiation of satellite cell activation, crucial steps that support the repair and remodeling of muscle fibers(48, 56). As shown in Figure 8, TNF*α* and IL1β, showed no significant difference between PNCOL (+) and PNCOL (-) groups. However, CD163 and CCL18 genes, associated with anti-inflammatory and tissue remodeling processes, were upregulated 1.5-fold and 3-fold, respectively in PNCOL (+) samples compared to the PNCOL (-) counterparts. group after the mechanical stimulation and PNCOL treatment at the damage site (Figure 8). These results indicate the potential efficacy of PNCOL in promoting muscle tissue repair and mitigating inflammation under mechanical loading in an ex-vivo organ culturing platform.

**Figure 8.**
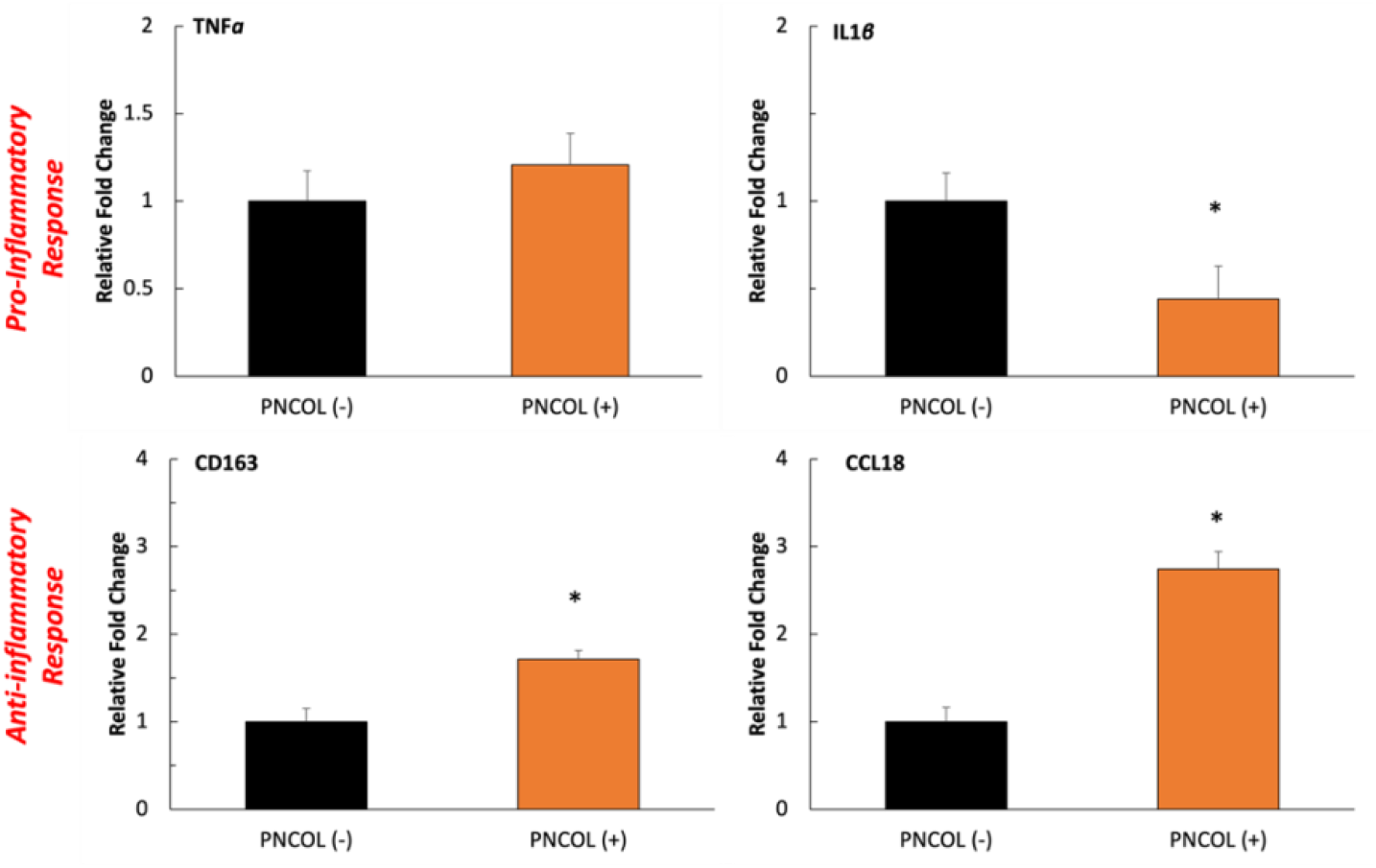
The relative fold changes of the inflammatory response of PNCOL (-) and PNCOL (+) samples following 7 days of organ culture in a dynamic mechanical environment. All data were expressed as the mean and standard errors. *Indicates the statistical differences between groups with p<0.05, n=4; at least three technical replicates were used for gene expression analysis.

### 2.7. Inflammation Stability of TA Muscle Defect Site After Treatments

Finally, to elucidate the impact of PNCOL treatment on TA defect under mechanical stimulation at the molecular level, protein expression via immunostaining was conducted. Figure 9 shows the presence of CD206 and *α*SMA surface markers in the PNCOL (+) compared to the PNCOL (-) group.

**Figure 9.**
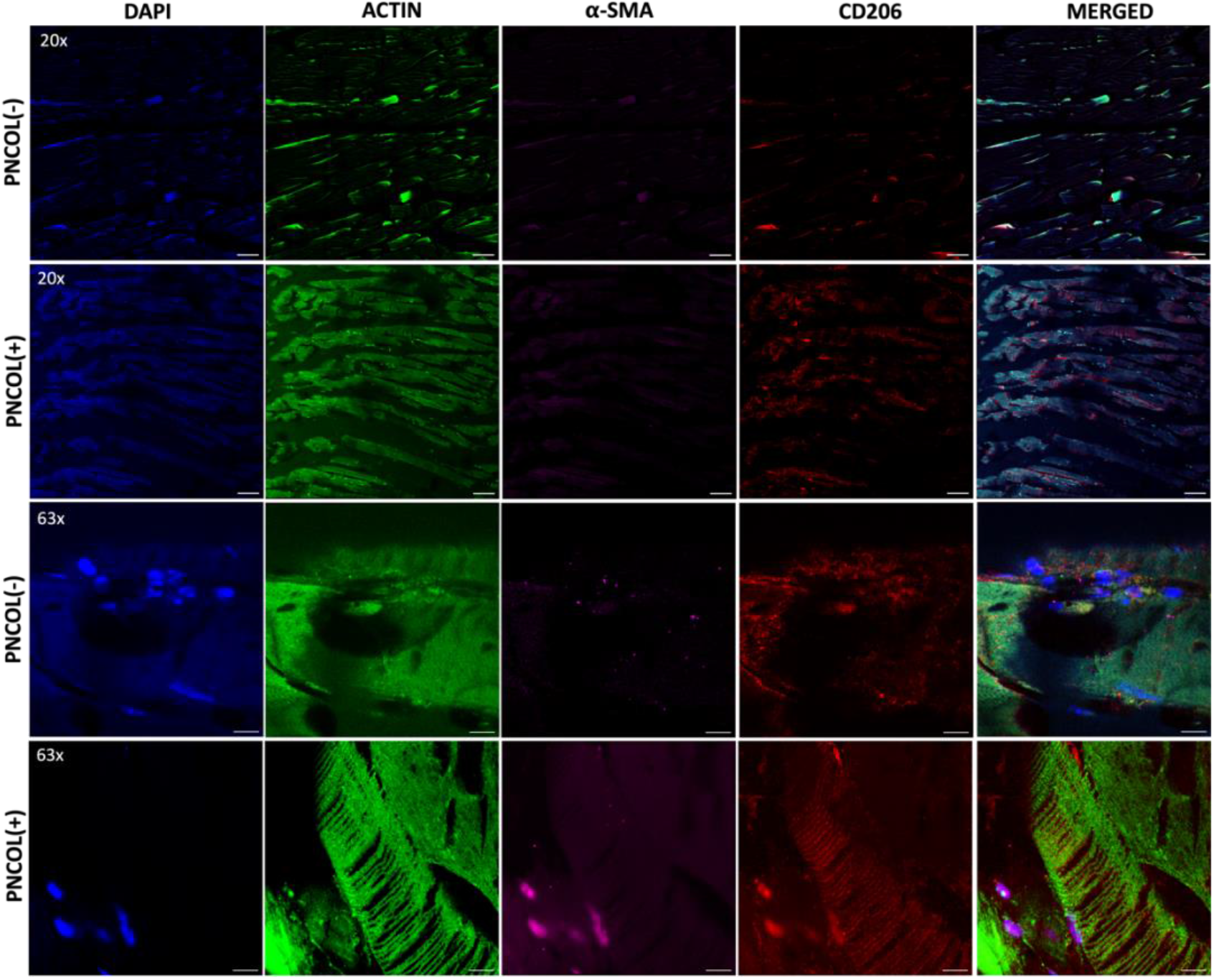
Antibody staining of PNCOL (-) and PNCOL (+) groups after 7 days of mechanical stimulation. Phalloidin (green) for f-actin staining, CD206 (red) as an anti-inflammatory marker, *α*SMA (magenta) for muscle regeneration marker, and DAPI (blue) for cell nuclei. *(n=4)*. 20x images, the scale bar represents 100 µm, and 63x images, the scale bar represents 10 µm.

The CD206, a marker associated with anti-inflammatory processes and tissue remodeling, and *α*SMA, a marker indicative of fibrotic response, reflect the modulation of immune responses and fibrotic reaction towards enhanced muscle regeneration in the PNCOL (+) group compared to the control. Since monitoring actin expression and localization can offer valuable information about the progression and efficacy of the regeneration process in musculoskeletal tissues (57-59), the changes in actin in PNCOL (+) samples were observed and compared with the PNCOL (-) group. Figure 9 demonstrates a more dominant presence of actin in the PNCOL (+) compared to the PNCOL (-) group as seen in all 20x figures and later confirmed in the zoom-in figures (63x). This relevant presence of actin in the PNCOL (+) group may indicate enhanced cytoskeletal rearrangement and cell migration, suggesting that PNCOL treatment could promote more efficient tissue repair processes through modulation of actin dynamics.

## 3. Discussion

Recognizing the significance of whole organ culture in replicating complex physiological environments is necessary for advancing our understanding of tissue response and enhancing better therapeutic outcomes. Thus, in this study, not only an innovative ex-vivo hindlimb dynamic organ culturing platform is introduced but also the role of injectable cell-laden nanofibrous matrix in critical size defect TA muscle regeneration was studied under physiologically relevant mechanical loading. Before extensive biological and structural analyses, we first aimed to demonstrate the transmission of mechanical loading applied to the organ throughout its structure, by characterizing the bone structure. In ex-vivo organ culturing, upon mechanical loading, notably, the cortical bone of the tibia and femur exhibited increases in total area and marrow area, without changes in cortical bone mass, which can be interpreted as a result of increased periosteal bone formation, where bone and muscle is juxtaposed, with simultaneously increased bone resorption at the endosteal surface which may indicate increased proinflammatory environment in an absence of blood perfusion and innervation. The concept of muscle and bone being intertwined in their functions and physiology suggests that changes in one tissue can significantly impact the other(60). These together with demonstrating that cell viability and hindlimb integrity are maintained throughout the experiment reassure the efficacy of the ex-vivo hindlimb organ culturing platform.

The optical images (days 0 and 7) of the hindlimbs cultured using a dynamic organ culture platform demonstrated the gross health of the samples throughout treatments (Figure 2A). The cell viability, assessed using the colorimetric-XTT assay, remained preserved throughout the 7-day organ culture period for both PNCOL (-) and PNCOL (+) samples (Figure 2B). Importantly, there was no statistical difference in cell viability between day 1 and day 7 for either group, suggesting that the ex-vivo mice hindlimb culturing platform, irrespective of PNCOL treatment, maintains consistent cell viability over the culturing period. Overall, the µ-CT images and image-based quantifications (Figure 1), the optical images, and cell viability data (Figure 2) suggest that mice hindlimb organ culturing platform under dynamic mechanical loading conditions can transmit the mechanical loading from muscle to the bone as experienced in locomotion while preserving the viability of the organ. Thus, the ex-vivo mice hindlimb organ culturing platform used in this study offers a valuable tool for investigating the isolated effects of variables such as mechanical loading, biomatrix for bone or muscle regeneration, electrical stimulation, ultrasound treatment, and more. Unlike in vivo studies, this platform provides a controlled environment that minimizes costs and enhances consistency. By isolating various factors that may influence outcomes, researchers can gain deeper insights into tissue responses without the complexities inherent in in vivo studies.

Once the viability of the ex-vivo mice hindlimb organ culturing platform was established, a detailed analysis was conducted to understand the potential of myoblast-laden injectable matrix (PNCOL) in TA muscle regeneration under dynamic loading of 12% with 1Hz frequency within the ex-vivo mice hindlimb organ culturing platform. The histological images of the damaged site (Figure 3) demonstrated that the TA defect treated with PNCOL had a high number of nuclei (purple dots within H&E images in Figure 3) corresponding to the presence of satellite cells and higher muscle regeneration which further was confirmed with dense G-Trichrome and Desmin presence in PNCOL (+) hindlimbs compared to the PNCOL (-) group. The PNCOL (+) group regenerated the muscle histomorphology at the defect site. The normalized proteoglycan content data (Figure 4) agreed with the histological data suggesting that GAG content increased statistically significantly more than 3-fold in TA treated with PNCOL (+) over the organ culturing period. Overall, the histological data along with the GAG content data suggested that there was a successful de novo extracellular matrix (ECM) regeneration following the injury for the TA muscle treated with the PNCOL. The newly generated ECM fibers at the defect site were visualized with SEM (Figure 5). The SEM micrographs showed that the gap at the defect site was woven with the ECM fibers in PNCOL (+) groups while there was a visible gap for PNCOL (-) group after 7 days.

Following structural and quantitative analyses of the ECM matrix component, the proteomic and gene expression analyses were conducted to understand the cellular and sub-cellular level changes leading to higher ECM production for the PNCOL (+) groups. The cytokine and chemokine production were analyzed through the media on individual ex-vivo organ cultures after 7 days. For instance, the higher expression of adiponectin in the PNCOL (+) group seen in the cytokine profile after the treatments might describe a favorable shift toward regeneration. The role of adiponectin appears to have a direct impact on muscle tissue regeneration(61). The cytokine profile demonstrated an increased expression of CXCL1, CXCL2, and CXCL16 chemokines regulating inflammation and immune cell recruitment, which are crucial processes in tissue repair and regeneration. These chemokines are implicated in orchestrating the early stages of neutrophil, macrophage, and other immune cell infiltration into injured muscle tissue, thereby facilitating muscle regeneration(62, 63). Also, insulin-like growth factor-binding proteins (IGFBPs) regulate bone metabolism and play a crucial role in modulating the bioavailability and activity of IGFs, which are regulators of bone growth and remodeling(64). IGFBP5 and IGFBP6 demonstrated a higher expression in the PNCOL (+) compared to the untreated counterparts. The intricate balance between IGFBPs influences bone formation and resorption processes during the crosstalk between muscle, bone, and endocrine factors in maintaining musculoskeletal health(64). Overall, the cytokine and chemokine profile array demonstrated a positive response to both PNCOL and mechanical stimulation in 7 days of ex-vivo organ culturing. Since the media extracted from each culture may have other components and smaller no-present proteins were not expressed on our proteome profile, gene expression analysis was conducted to thoroughly identify the regenerative capabilities of the bone-muscle organ after the treatments.

The relative gene expression of α-SMA, PAX7, and Myogenin genes was increased significantly in the PNCOL (+) group in comparison to the PNCOL (-) group. α-SMA is an actin expressed by myofibroblasts, and the increased presence of αSMA denotes that myofibroblasts incorporated with the PNCOL have survived and proliferated (57, 65). Similarly, MYOD is a vital regulator of skeletal muscle metabolism, and no significant difference in expression was found between the PNCOL (+) and PNCOL (-) groups. In tandem with myofibroblasts’ increased survival and proliferation, the stagnating MYOD expression is consistent. Muscle cell metabolism regulation remaining unchanged for a larger population of cells allows more cells to survive (66). The PAX7 gene is uniformly expressed by satellite cells in human muscle tissue and is a significant component in the regenerative properties of muscle tissue (50, 67). Moreover, Figure 7 contains the gene expression of Mrf4, MYF5, and TGFB1 which significantly increased expression in the PNCOL (+) group while VEGFA remained constant. Mrf4 functions to negatively regulate MSK growth, whereas MYF5 functions to promote growth (68). These two genes enforce homeostasis between the promotion and restriction of muscle growth. Though increased expression of Mrf4 alludes to hindered muscle growth, the greater expression of protein MYF5 relative to the increase in Mrf4 shows the presence of increased regeneration. TGFB1 belongs to the protein family of transformative growth factors and promotes the proliferation of MSK cells (69). Additionally, VEGFA functions as a regulator for various endothelial cell activities such as proliferation and migration. Though endothelial tissue is irrelevant in this study, the role of VEGFA is to act as a diagnostic for the overall health of the tissue. The increase of Mrf4, MYF5, and TGFB1 supports the idea of bolstered muscle regeneration within the PNCOL (+) group (70, 71). A similar expression of VEGFA between the two groups suggests the continued health of the tissue.

Figure 8 demonstrates TNF*α*, IL1β, CD163, and CCL18 gene expression. TNF*α* functions as a regulator of the inflammatory response and expressed no statistical difference between the PNCOL (+) and PNCOL (-) groups. Instead, a minor increase in expression in the PNCOL (+) group was seen; this is likely the product of the increased presence of muscle regeneration factors in the PNCOL (+) group. IL1β is a cytokine interleukin and is integral to the tissue-bound inflammatory response(72). Most notably, IL1β exacerbates cell-wide damage during the inflammatory response greatly, yet not the regenerative factors associated with the inflammation(73). CD163 is a macrophage-based protein and is an additive marker for tissues responding to inflammation (74). CCL18 is a scientifically ambiguous protein and may have functions of chronic inflammation (75). The fluctuations of protein expressions across the PNCOL (+) and PNCOL (-) support the idea of PNCOL causing an overall larger inflammation response than the control. CCL18 is expressed at a much more significant fold change, and though IL1β and CD163 present an inconsistency, the respective decreases and increases of each protein are not significant relative to the increase of CCL18.

This suggestion is further supported by immunostaining qualitative analysis (Figure 9) with increased tissue regeneration and the presence of anti-inflammatory markers. Qualitative evidence from histology and histomorphometry display long, smooth bands of muscle cells in the presence of collagen, suggesting heightened MSK regeneration in the PNCOL (+) group. Overall, the data demonstrates that the ex-vivo organ culturing platform can keep the bone and muscle tissue functions over 7 days of the culturing period under mechanical loading conditions. In addition, the data proves that the critical size defect in TA muscle can be treated with PNCOL injection to the defect site. Our study underscores the efficacy of PNCOL treatment combined with 3D mechanical loading using an ex-vivo organ culturing platform for promoting muscle tissue regeneration and reducing inflammation. These findings demonstrate the importance of multidimensional approaches for enhancing therapeutic outcomes in MSK disorders.

In addition, drawing on recent debates in the field of bioethics, this study also develops an interdisciplinary approach and adheres to the 4Rs principle (Reduction, Refinement, Replacement and Responsibility) for ethical animal research (76, 77) through employing an ex-vivo approach with isolated mouse hindlimbs instead of in-vivo testing. Our motivation aligns with the growing emphasis on minimizing the number of animals used in research (78) and developing a robust sense of responsible animal experimentation.

It is important to note that while the ex-vivo hindlimb dynamic organ culturing platform along with cell-laden injectable PNCOL matrix offers valuable insights into muscle tissue regeneration, this study has some limitations as well. For instance, its scalability to larger animals may be limited. This platform may primarily be applicable to mice and rats, which exhibit notable differences in muscle regeneration compared to humans. Furthermore, the use of the C2C12 myoblast cell line, while informative, may not fully represent the success/ effects of delivering primary myoblast cultures directly from mouse skeletal muscle tissue. Additionally, the fate of the delivered cells remains unclear, warranting further investigation in future studies. Future studies should also consider employing additional markers or techniques, such as immunohistochemistry or single-cell RNA sequencing, to further characterize the specific cell populations at the defect site to distinguish the mixture of different cell types, including satellite cells, fibroblasts, macrophages, and myoblasts.

## 4. Methods and Materials

### 4.1. Ex-Vivo Organ Culture Platform and Mouse Hindlimb Culturing under Mechanical Loading

The organ culture platform, an innovative, custom-built mechanical loading platform, was designed to create physiologically relevant mechanical stimuli for the whole bone-muscle organ. Unlike other mechanical loading platforms, our proposed design creates predefined uniform mechanical strain on the whole organ while maintaining the stability of the tissues. The organ culture platform (Figure 10) is designed to load the entire bone-muscle organ onto the platform through fixed grips at the top and bottom ends. The uniaxial tension of the organ produces mechanical strain in the bone and muscle tissues. This gives us control over the movement of the entire organ structure and control over the amount and duration of strain applied, simulating different physiologically relevant environments. The uniaxial tensile organ culture platform’s working mechanism detailed in this protocol, provides mechanical stimulation through a stepper motor displacement. This type of stimuli has been widely used to simulate heart muscle and musculoskeletal tissues, including tendon, ligament, bone, and cartilage, through cyclic tensile force(79-81). Figure 10 shows the schematic of the ex-vivo hindlimb organ culturing platform with mechanical loading.

**Figure 10.**
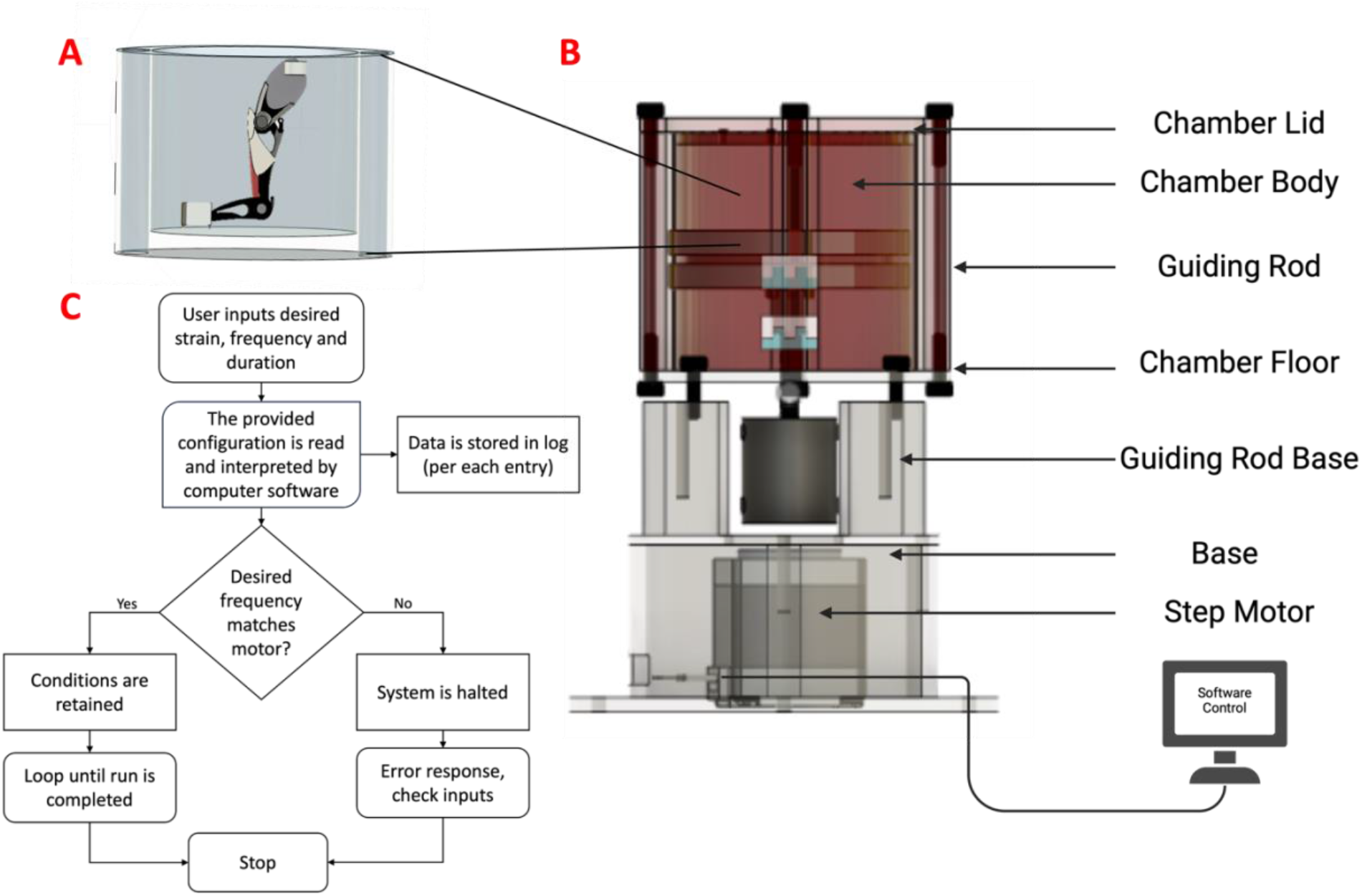
Organ Culture Platform Components and System Flowchart. **A)** The schematic representation of mouse hindlimb within the biocompatible organ culturing chamber. **B)** The components of the organ culturing platform. **C)** Architectural and Functional schematic of hardware and software of the organ culture platform.

As shown in Figure 10A, the chamber is an elliptical cylinder comprised of biocompatible resin, a biologically compatible material (Formlabs, Somerville, U.S.). The chamber lid and chamber floor seal the longitudinal ends of the chamber. The chamber is restricted in axial movement using guide rods. The guiding rods are affixed to the guiding rod base via threading and stainless-steel screws, and the guiding rod base is affixed to the base identically. The aperture in the guiding rod base allows for the protrusion and retraction of the stepper motor, which moves vertically and provides tensile stress to the legs in the chamber. Through a wiring system, the organ culture platform is connected to a computer interface to control the platform. The sample is attached via stainless steel clamps and an adhesive from the lid to the floor of the organ culture platform (Figure 10A). The functional map shown below in Figure 10C provides the hardware and software logic of the organ culture platform.

#### Ex-vivo Mouse Hindlimb Culturing

To test the efficacy of the organ culturing platform, ex-vivo mouse hindlimbs culturing was conducted under physiologically relevant mechanical stimulation for 2 hours a day for 7 days. The ex vivo mouse hindlimbs were subjected to 12% uniaxial tensile strain at a frequency of 0.1 Hz to replicate the dynamic physiological conditions of MSK tissues, including tendon, ligament, and muscle. The 12% uniaxial strain corresponds to the stress encountered by skeletal tissues, mirroring the strain experienced during exercise, which can reach up to 12.5 times body weight. This strain also reflects the length changes of 5-9% observed during moderate physical activities and up to 12% during high-intensity exercise (57).

For ex-vivo mouse hindlimbs culturing, the 8-month-old, wild-type male mice (strain C57BL/6) were used. These relatively young, physically mature male mice are analogous to young, physically fit athletes, susceptible to muscle injury(82-84). All dissection procedures followed protocols approved by the Institutional Animal Care and Use Committee and the University of Toledo’s approval. Briefly, mice were euthanized using CO_2_ in a sterile environment. The abdomen was then cut and separated using blunt-end scissors, and surface muscles were removed to expose the pelvic-hip joint. The pelvic hip joints of both hindlimbs were then cut, and the hindlimbs were removed from the abdomen. The skin and hair of each hindlimb were removed with blunt dissection. Then, hindlimbs were placed on ice in phosphate-buffered saline solution (PBS) for 2 hours. The hindlimbs were then placed into an organ culture platform with DMEM media supplemented with FBS 10%, 1% P/S, 1:500 Primocin, 1.5% 0.4 M NaCl, and 1.5% 0.4 M KCl for culturing for 7 days.

### 4.2. Micro-CT Analysis of Mouse Hindlimb after Ex-Vivo Organ Culturing under Mechanical Loading

Assessment of cortical bone in the tibia and femur was conducted by micro-CT using the µCT 35 system (Scanco Medical AG, Bruettisellen, Switzerland). Bone scans were performed with the x-ray source operating at 70 kVp and 114 µA energy settings and recording 500 projections/180° acquired at 300 ms integration time using 7 µm nominal resolution voxel for both bone locations. Scans of the cortical bone at the tibia and femur midshaft contained 57 slices all of which were contoured automatically, and which were segmented at 260 per mille threshold. Simulated bone bending strength (I_max_/C_max_ and I_min_/C_min_) and torsional strength (pMOI) at the tibia midshaft were based on bone cross-sectional geometry in combination with local tissue mineral content(85). The analysis of the trabecular bone microstructure, cortical bone parameters, and simulated bone strength was conducted using Evaluation Program V6.5-3 (Scanco Medical AG) and conformed to recommended guidelines(86).

### 4.3. Myoblast Cell-laden Injectable Matrix Synthesis for TA Muscle Regeneration

#### 4.3.1. Cell-laden Injectable Nanofibrous Matrix for Tibialis Anterior Muscle Regeneration

Cell-based injectable tissue scaffolds play a pivotal role in tissue regeneration by providing a supportive framework for cellular growth and organization for restoring damaged tissues. To regenerate damaged Tibialis Anterior muscle, an injectable and nanofibrous called PNCOL matrix was synthesized based on our well-established protocols (87-89). Figure 11 demonstrates the schematic representation of the cell-laden injectable PNCOL matrix synthesis process.

**Figure 11.**
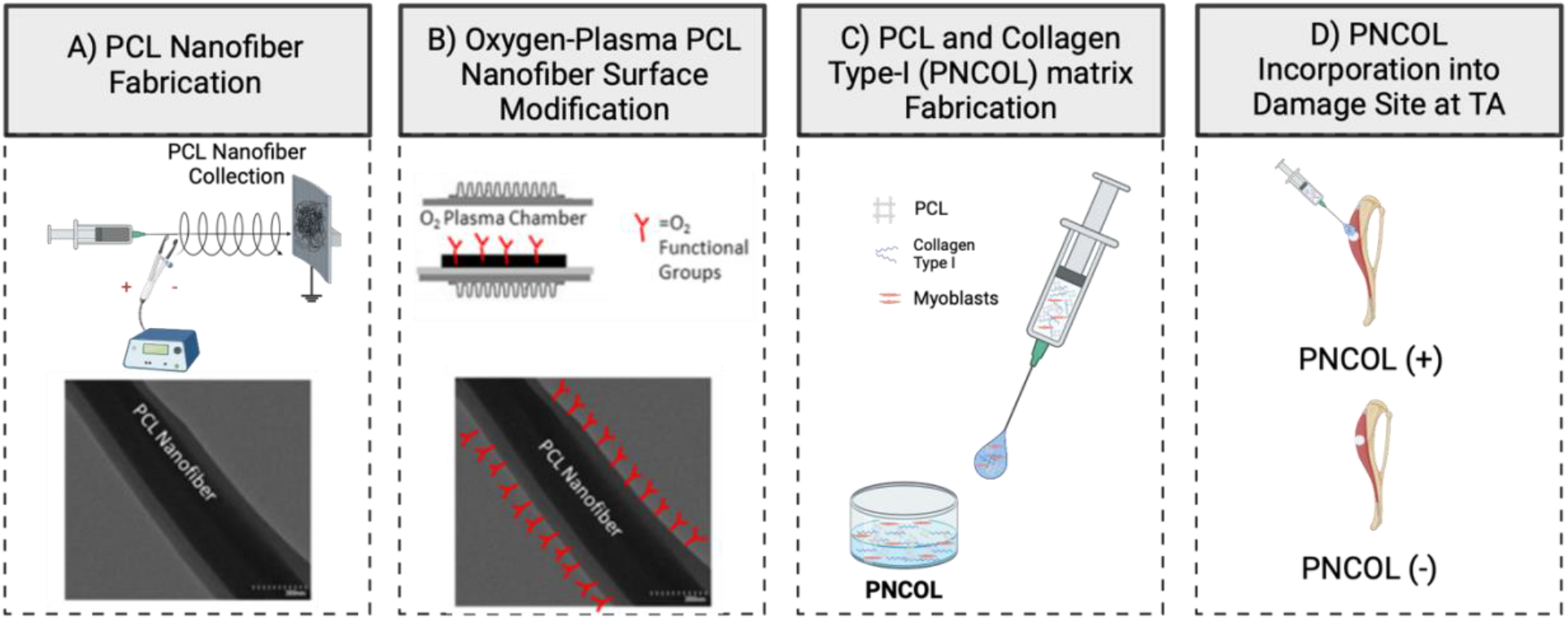
Myoblast cell-laden PNCOL (polycaprolactone and collagen type-I) matrix synthesis to be injected at Tibialis Anterior (TA) Muscle defect.

Briefly, PCL (MW 45,000, Sigma-Aldrich, St. Louis, MI, USA) pellets were dissolved overnight in a 3:1 mixture of chloroform and methanol solution under a sterilized chemical hood at 16% w/v. The following day, the solution was loaded into a syringe pump and extruded with a flow rate of 8 mL/hour, through a 20-gauge needle. A 20 kV potential was applied between the needle tip and a collector plate where PCL nanofibers were collected (Figure 11A). Following 2 hours of PCL nanofiber collection, the PCL fibers were transferred under the chemical hood for residual solvent evaporation overnight. The following day, the PCL nanofibers were homogenized using a high-speed homogenizer (Ultra-Turrax, IKA Works, Inc., Staufen, Germany) and functionalized using oxygen-plasma treatment for 3 min to reduce its hydrophobicity based on our prior studies (87-89). Upon incubation, the PCL nanofibers were mixed within the neutralized collagen type-I solution with a 3% (w/v) concentration (Figure 11B-C).

The neutralized collagen type-I solution with 3 mg/mL concentration was prepared from 9.1 mg/mL collagen type-I stock solution (Corning, Corning, NY, USA) with a pH of ∼3.4 using 1 M NaOH, phosphate-buffered solution (PBS) and culture media. Then, the mouse myoblasts cell line (C2C12, ATCC, USA) cultured in a complete media of Dulbecco’s Modified Eagle’s medium (DMEM) (ATCC; Manassas, VA, USA) supplemented with 10% fetal bovine serum (FBS) (Corning; USA) and 1% penicillin-streptomycin (Corning; USA) was encapsulated within the PNCOL with a cell density of 10^6^ cells/mL (Figure 11C).

#### 4.3.2. TA Muscle Defect Creation and Cell-Laden Matrix Injection to the Defect Site

Following extraction, critical size creation of muscle loss injury was performed on the tibialis anterior (TA) muscle using a 3 mm biopsy punch (Robbins Instruments Chatham, NJ, USA) on each ex-vivo mouse hindlimb. TA muscle corresponds to 0.17% of the total body weight(6, 24), excision of approx. 20% of the TA muscle weight in the middle third of the sample characterized a critical size damage injury. Following defect creation, 200 uL of the myoblast-laden nanofibrous matrix, called PNCOL, was injected into the defect areas. Hindlimbs in which the defect site received no PNCOL treatment served as a control. Then, all defect sites were sutured using the modified purse-string suture techniques (MPSS) based on our established protocol(90). Two overlapping suture loops were interconnected and contracted circumferentially for each defect site to provide near-watertight sealing. Following suturing, all sample groups were moved to the organ culture chamber for 7 days under 2 hours of mechanical loading as explained in Section 2.1.

### 4.4. Assessing the Effect of Cell-laden Injectable Matrix for TA Muscle Regeneration Following Ex-vivo Organ Culture under Dynamic Mechanical Loading

The TA muscle regeneration was characterized using structural and biological assessment including gross defect morphology, histology, and antibody staining. Defect morphology was measured by photographing and weighing the experimental and control groups before and after the 7 days. Post healing, 5 μm long longitudinal samples were retrieved roughly 400μm deep in the damaged area. These samples were used in concurrence with Hematoxylin & Eosin and Masson’s trichrome stains to identify inflammation around the PNCOL injected area (damage site). Gene expression, cytokine production, and protein analysis were utilized to assess the regenerative effect of the treatments at the damage site (TA muscle). Figure 12 summarizes the design and characterization of this section of the study.

**Figure 12.**
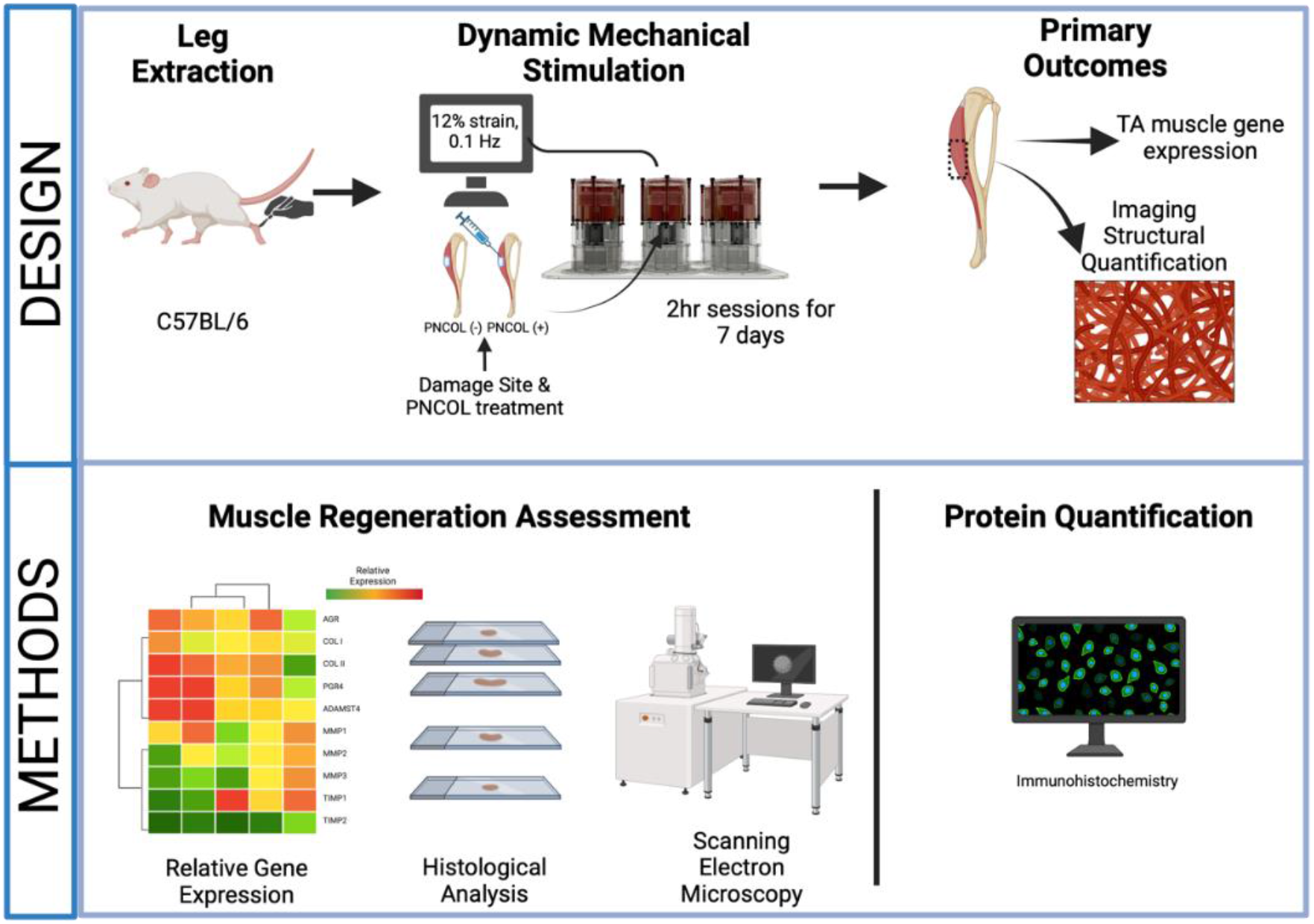
The major steps followed in the study.

#### 4.4.1. Structural and Morphological Characterizations of the Ex-vivo Mouse Hindlimb Organ Culturing under Mechanical Loading

##### Scanning Electron Microscopy Analysis

TA muscle, PNCOL matrix, and collagen fiber morphology were characterized using Scanning Electron Microscopy (SEM) (Hitachi, Santa Clara, CA, USA) after ex-vivo organ culturing with mechanical stimulation. Before characterization, TA from the damaged site were cut and fixed with 4% paraformaldehyde in PBS for 30 min. After fixation, the samples were dehydrated first in sequential ethanol solutions with increasing concentrations from 30% to 100% for 15 min each. The samples were then further dehydrated by being submerged into sequential ethanol/hexamethyldisilane (HMDS) solutions from 30% to 100% for 10 minutes each to improve optical image quality. After the chemical treatment, the samples were left to air-dry overnight. The gold-sputter coating was applied to the samples the following day, allowing for visualization under SEM to observe structural and morphological changes upon experimental setup.

##### Histological Analysis

Variations in cell density, collagen structure, and tissue inflammation were analyzed using Histological staining. Muscle tissue samples extracted from the damage site of PNCOL (-) and PNCOL (+) groups were submerged in 10% normalized formalin for 48 hours. The tissue samples were then processed on a Sakura Tissue TEK VIP 5 tissue processor, dehydrating the samples in 70%, 85%, 95%, and 100% ethanol before being treated in a clearing agent (FISHER XS-05 laboratory-grade Xylene). After dehydration treatment, samples were sealed in paraffin (Leica Paraplast Plus) and implanted into a tissue cassette before being incised on a LEICA RM 2235 microtome in 5 mm sections and placed on slides. The slides were incubated in hematoxylin and eosin (H&E), and Masson’s trichrome staining, washed, and viewed under a bright field microscope.

#### 4.4.2. Biological Characterization of the Ex-vivo Mouse Hindlimb Organ Culturing under Mechanical Loading

##### Cell Viability Analysis

Cell viability on the mouse hindlimb culture was assessed on days 1 and 7 using the XTT cell viability assay kit (Sigma Millipore, Product No. 11465015001, USA). This assay measures metabolic activity as an indicator of cell viability, specifically focusing on mitochondrial enzyme activity. Formazan production, directly proportional to the number of metabolically active cells, occurs when XTT is reduced in viable cell mitochondria. The absorbance of the resulting-colored solution correlates with cell viability, determined by comparing samples. Triplicates of 100 μL of media extracted from each organ culture chamber were incubated on a 96-well plate and mixed with 70 μL of XTT assay reagent. Following an 8-hour incubation at 37°C, the wells’ optical density (OD) was measured using a UV kinetic microplate reader at 450 nm, with a reference wavelength of 660 nm. Each plate included appropriate blank control wells containing media without reagent. Absorbance readings were calculated according to the vendor’s protocol and extrapolated to the initial time point.

##### Proteome Array Analysis

The Proteome Profiler Mouse XL Cytokine Array kit (R&D Systems, Minneapolis) was used to analyze cytokine, chemokine, and growth factors released from whole hindlimbs in culture after 7 days of mechanical stimulation and PNCOL treatment. Each kit provided four membranes, sufficient for four distinct samples. The membranes featured 111 capture antibodies in duplicate, targeting various biomolecules as listed on the manufacturer’s website. Conditioned media samples (1-2 mL) were collected and stored on ice during collection after 7 days of treatments (mechanical stimulation and PNCOL), then stored at -80°C until further use. Upon thawing, samples underwent centrifugation at 2,000 rpm for 3 minutes to remove cellular debris. The supernatant, devoid of cellular components, was reserved for experimentation and maintained on ice throughout the process—experimental procedures adhered to the protocol outlined by the manufacturer. Image analysis was conducted using HLImage++ software (Western Vision Software) to interpret results. Cytokines with pixel density (PD) differences greater than 200 pixels between groups were selected. This threshold method facilitates the criteria aimed at identifying biomarkers indicative of significant changes in expression levels.

##### Gene Expression Analysis

The TA muscles (damage site) from PNCOL (+) and PNCOL (-) groups were extracted and dissected after experiment termination for gene expression analysis. Gene expression was carried out using real-time polymerase chain reaction (qRT-PCR). Tissue samples were extracted one day after the experiment’s termination and prepared by snap freezing and mechanical disruption. RNA was isolated using a Directo-zol RNA Extraction Kit (Zymo Research, USA). Per the manufacturer’s instructions, this isolated RNA was reverse transcribed using the Superscript IV Reverse Transcriptase (ThermoFisher, USA). Quantitative real-time PCR was performed using the iTaq Universal SYBR Green Supermix (Bio-Rad, Hercules, CA, USA) to investigate the expression of muscle regeneration markers and immune response markers. The relative gene expression for fold changes between PNCOL (-) and PNCOL (+) samples was obtained using the ΔΔCt method. The ΔΔCt method utilizes glyceraldehyde-3-phosphate dehydrogenase (GAPDH) as a housekeeping standardizing gene. RT-qPCR was performed in the iCycler iQ detection system (Bio-Rad, Hercules, CA, USA), with thermocycling for 35 cycles. The primer sequences were obtained from published literature and were designed and purchased from Integrated DNA Technologies (IDT, Coralville, IA, USA). The list of genes and associated primer sequences are listed in Table 1 in the Supplementary Document.

##### Immunostaining and Immunohistochemical Analysis

Building upon the gene expression analysis, further evaluation of the immune response following treatments involved assessing the TA muscle tissue at the damage site through immunostaining. Paraffin-embedded samples were sectioned into ∼40um thick slices using a microtome (GMI-Reichert Jung 820 II), mounted on glass slides, and cleared using a xylene/ethanol rehydration protocol. Following deparaffinization and rehydration, the slides were incubated in trypsin and calcium chloride solution (0.1% (v/v) trypsin (ATCC, USA) plus 0.1% (w/v) calcium chloride (Sigma-Aldrich, USA)) at 37°C for 30 min. Then, heat-mediated antigen retrieval was performed by immersing the slides in citrate buffer (10 nM sodium citrate in 0.02% Tween 20) at 95°C for 30 min. For immunostaining, samples were permeabilized with 0.1% Triton x-100 in PBS. Then, sections were washed and blocked in 2.5% goat serum and 0.2% Tween in PBS for 20min. After blocking, slides were incubated overnight at 4°C in mouse anti-Smooth Muscle Actin (1:40) (AB262054, Santa Cruz) and recombinant rabbit CD206 antibody (1:40) (ThermoFisher, MA1-35936). The next day, slides were washed with PBS and 0.2% Tween solution and blocked in 2.5% (v/v) goat serum for 20min. Then, slides were incubated with Alexa Four 488 Phalloidin (1:200) (ThermoFisher, A12379), goat anti-rabbit IgG (H+L) Highly Cross-Absorbed Secondary Antibody Alexa Flour 594 (1:200) (ThermoFisher, A32740) and Goat anti-Mouse IgG (H+L) Highly Cross-Absorbed Secondary Antibody, Alexa Flour 657 (ThermoFisher, A32728TR) for 2 hours. Finally, slides were incubated with DAPI (4’,6-Diamidino-2-Phenylindole) (Life Technologies, USA) dye (1:1000) in PBS for 30 min, washed with 0.2% tween. Slides were then imaged using a HC PL APO 63X/1.40 OIL CS2 or HC PL FLUOTAR L 20× air objective with a Leica Stellaris 5 confocal system equipped with HyD detectors and the LASX software. Confocal images from at least three independent studies were processed to visualize each cell’s nuclei, filamentous actin (F-actin), *α*SMA, and CD206 expressions. For desmin immunohistochemical staining, tissue sections after deparaffinization, heat-induced antigen retrieval, and non-specific binding sites blocked, were incubated with a primary antibody specific to desmin (1:200) (ThermoFisher, MA1-06401). After antibody incubation, the slides were washed. The target antigen was visualized using a chromogenic substrate specific to desmin, forming a brown precipitate at the antigen binding site. Slides were then imaged using a bright field microscope.

### 4.5. Statistical Analysis

Statistical analysis was executed through R-studio using the Student’s T-test. All values are reported as the mean ± the standard error. p < 0.05 represents a statistical difference. Symbols located on top of each bar profile condense the statistical analysis results. Each study of 2 samples was repeated twice (n=4). Two legs were assigned as a PNCOL-treated group and another two as the control group lacking any PNCOL treatment. At least three technical replicates were produced for each assay performed in this study. (*) Indicates the statistical differences between control and experimental groups with p<0.05, (#) indicates a statistical difference between experimental groups of the same day data point.

## 5. Conclusion

The novelty of this study arises from its combination of mechanical stimulation and injectable nanofibrous matrix scaffold treatment, PNCOL, onto a target damage site on an ex vivo mice hindlimb organ culture towards muscle regeneration. Notably, our experimental setup allowed us to simulate physiological loading observed in musculoskeletal tissues, providing a unique opportunity to investigate the broader implications of mechanical stimulation on the whole organ composition within an ex vivo culture setting.

## Author Contributions

Conceptualization, E.Y.-A., B. L.-C, and D.J.; Methodology, D.J., J.H., and E.Y.-A.; Formal analysis, D.J. and E.Y.-A; Investigation, D.J., J.H., E.C., A.R., R.G.-M., and E.Y.-A.; Writing—original draft, D.J., and E.Y.-A.; Writing—review & editing, J.H., B.L-C., P.C., A.R., M.Y, and R.G.-M.; Visualization, D.J., E.C., and A.R. All authors have read and agreed to the published version of the manuscript.

## Data Availability Statement

The document contains the data; otherwise, they can be shared by the corresponding author upon request.

## Abbreviations

CCL-18: Chemokine (C-C motif) ligand-18
CD163: Cluster of Differentiation 163
CD206: Cluster of Differentiation 206
IFN-*γ*: Interferon gamma
ECM: Extracellular matrix
FBS: Fetal bovine serum
GAPDH: Glyceraldehyde-3-phosphate dehydrogenase
IL: Interleukin
IL1β: Interleukin beta 1
LPS: Lipid polysaccharide
VEGFA: Vascular endothelial growth factor A
TGFB1: Transforming growth factor beta 1
ECM: Extracellular Matrix
MMP: Matrix metalloproteinase
MRF4: Myogenic regulatory factor 4
MYF5: Myogenic regulatory factor 5
PBS: Phosphate-buffered saline
PD: Pixel Density
PCR: Polymerase chain reaction
PNCOL: Polycaprolactone nanofiber and collagen
qRT-PCR: Quantitative real-time polymerase chain reaction
RNA: Ribonucleic acid
RPMI: Roswell Park Memorial Institute
TNF-α: Tumor necrosis factor-α
C2C12: Immortalized mouse myoblast cell line

## List of Units and Symbols

M: Molar
U: Unit
v/v: Volume per volume
w/v: Weight per volume
μ: Micro

## References

1. Lloyd D. The future of in-field sports biomechanics: wearables plus modelling compute real-time in vivo tissue loading to prevent and repair musculoskeletal injuries. Sports Biomech. 2021:1–29. Epub 20210908. doi: 10.1080/14763141.2021.1959947. PubMed PMID: 34496728.

2. Thompson WR, Scott A, Loghmani MT, Ward SR, Warden SJ. Understanding Mechanobiology: Physical Therapists as a Force in Mechanotherapy and Musculoskeletal Regenerative Rehabilitation. Phys Ther. 2016;96(4):560–9. Epub 2015/12/08. doi: 10.2522/ptj.20150224. PubMed PMID: 26637643; PMCID: PMC4817213.

3. Yan Z, Yin H, Nerlich M, Pfeifer CG, Docheva D. Boosting tendon repair: interplay of cells, growth factors and scaffold-free and gel-based carriers. J Exp Orthop. 2018;5(1):1. Epub 20180105. doi: 10.1186/s40634-017-0117-1. PubMed PMID: 29330711; PMCID: PMC5768579.

4. Zhang Q, Adam NC, Hosseini Nasab SH, Taylor WR, Smith CR. Techniques for In Vivo Measurement of Ligament and Tendon Strain: A Review. Ann Biomed Eng. 2021;49(1):7–28. Epub 20201006. doi: 10.1007/s10439-020-02635-5. PubMed PMID: 33025317; PMCID: PMC7773624.

5. Huang C, Holfeld J, Schaden W, Orgill D, Ogawa R. Mechanotherapy: revisiting physical therapy and recruiting mechanobiology for a new era in medicine. Trends Mol Med. 2013;19(9):555–64. Epub 20130618. doi: 10.1016/j.molmed.2013.05.005. PubMed PMID: 23790684.

6. Lee EM, Kim AY, Lee EJ, Park JK, Lee MM, Hwang M, Kim CY, Kim SY, Jeong KS. Therapeutic effects of mouse adipose-derived stem cells and losartan in the skeletal muscle of injured mdx mice. Cell Transplant. 2015;24(5):939–53. Epub 20140303. doi: 10.3727/096368914×678599. PubMed PMID: 24593934.

7. Mouly V, Aamiri A, Bigot A, Cooper RN, Di Donna S, Furling D, Gidaro T, Jacquemin V, Mamchaoui K, Negroni E, Perie S, Renault V, Silva-Barbosa SD, Butler-Browne GS. The mitotic clock in skeletal muscle regeneration, disease and cell mediated gene therapy. Acta Physiol Scand. 2005;184(1):3 –15. doi: 10.1111/j.1365-201X.2005.01417.x. PubMed PMID: 15847639.

8. Qazi TH, Duda GN, Ort MJ, Perka C, Geissler S, Winkler T. Cell therapy to improve regeneration of skeletal muscle injuries. J Cachexia Sarcopenia Muscle. 2019;10(3):501–16. Epub 20190306. doi: 10.1002/jcsm.12416. PubMed PMID: 30843380; PMCID: PMC6596399.

9. Zhang Z, Zhao X, Wang C, Huang Y, Han Y, Guo B. Injectable conductive micro-cryogel as a muscle stem cell carrier improves myogenic proliferation, differentiation and in situ skeletal muscle regeneration. Acta Biomater. 2022;151:197–209. Epub 20220821. doi: 10.1016/j.actbio.2022.08.036. PubMed PMID: 36002125.

10. Kesireddy V. Evaluation of adipose-derived stem cells for tissue-engineered muscle repair construct-mediated repair of a murine model of volumetric muscle loss injury. Int J Nanomedicine. 2016;11:1461 –73. Epub 20160408. doi: 10.2147/IJN.S101955. PubMed PMID: 27114706; PMCID: PMC4833361.

11. Anloague A, Mahoney A, Ogunbekun O, Hiland TA, Thompson WR, Larsen B, Loghmani MT, Hum JM, Lowery JW. Mechanical stimulation of human dermal fibroblasts regulates pro-inflammatory cytokines: potential insight into soft tissue manual therapies. BMC Res Notes. 2020;13(1):400. Epub 2020/08/29. doi: 10.1186/s13104-020-05249-1. PubMed PMID: 32854782; PMCID: PMC7457292.

12. Wufuer M, Lee G, Hur W, Jeon B, Kim BJ, Choi TH, Lee S. Skin-on-a-chip model simulating inflammation, edema and drug-based treatment. Sci Rep. 2016;6:37471. Epub 2016/11/22. doi: 10.1038/srep37471. PubMed PMID: 27869150; PMCID: PMC5116589.

13. Schemitsch EH. Size Matters: Defining Critical in Bone Defect Size! J Orthop Trauma. 2017;31 Suppl 5:S20–S2. doi: 10.1097/BOT.0000000000000978. PubMed PMID: 28938386.

14. Shayan M, Huang NF. Pre-Clinical Cell Therapeutic Approaches for Repair of Volumetric Muscle Loss. Bioengineering (Basel). 2020;7(3). Epub 20200820. doi: 10.3390/bioengineering7030097. PubMed PMID: 32825213; PMCID: PMC7552602.

15. Bialorucki C, Subramanian G, Elsaadany M, Yildirim-Ayan E. In situ osteoblast mineralization mediates post-injection mechanical properties of osteoconductive material. J Mech Behav Biomed Mater. 2014;38:143–53. Epub 2014/07/23. doi: 10.1016/j.jmbbm.2014.06.018. PubMed PMID: 25051152.

16. Nasir NJN, Arifin N, Noordin K, Yusop N. Bone repair and key signalling pathways for cell-based bone regenerative therapy: A review. J Taibah Univ Med Sci. 2023;18(6):1350–63. Epub 20230524. doi: 10.1016/j.jtumed.2023.05.015. PubMed PMID: 37305024; PMCID: PMC10248876.

17. Ukeba D, Sudo H, Tsujimoto T, Ura K, Yamada K, Iwasaki N. Bone marrow mesenchymal stem cells combined with ultra-purified alginate gel as a regenerative therapeutic strategy after discectomy for degenerated intervertebral discs. EBioMedicine. 2020;53:102698. Epub 20200304. doi: 10.1016/j.ebiom.2020.102698. PubMed PMID: 32143180; PMCID: PMC7057222.

18. Fleming JW, Capel AJ, Rimington RP, Player DJ, Stolzing A, Lewis MP. Functional regeneration of tissue engineered skeletal muscle in vitro is dependent on the inclusion of basement membrane proteins. Cytoskeleton (Hoboken). 2019;76(6):371–82. Epub 20190819. doi: 10.1002/cm.21553. PubMed PMID: 31376315; PMCID: PMC6790946.

19. Ragini B, Kandhasamy S, Jacob JP, Vijayakumar S. Synthesis and in vitro characteristics of biogenic-derived hydroxyapatite for bone remodeling applications. Bioprocess Biosyst Eng. 2024;47(1):23 –37. Epub 20231112. doi: 10.1007/s00449-023-02940-y. PubMed PMID: 37952238.

20. Gharibshahian M, Alizadeh M, Kamalabadi Farahani M, Salehi M. Fabrication of Rosuvastatin-Incorporated Polycaprolactone -Gelatin Scaffold for Bone Repair: A Preliminary In Vitro Study. Cell J. 2024;26(1):70–80. Epub 20240131. doi: 10.22074/cellj.2023.2009047.1391. PubMed PMID: 38351731; PMCID: PMC10864776.

21. Akhter MN, Hara ES, Kadoya K, Okada M, Matsumoto T. Cellular Fragments as Biomaterial for Rapid In Vitro Bone-Like Tissue Synthesis. Int J Mol Sci. 2020;21(15). Epub 20200727. doi: 10.3390/ijms21155327. PubMed PMID: 32727114; PMCID: PMC7432235.

22. Lawrence MM, Van Pelt DW, Confides AL, Hettinger ZR, Hunt ER, Reid JJ, Laurin JL, Peelor FF, 3rd, Butterfield TA, Miller BF, Dupont-Versteegden EE. Muscle from aged rats is resistant to mechanotherapy during atrophy and reloading. Geroscience. 2021;43(1):65–83. Epub 20200625. doi: 10.1007/s11357-020-00215-y. PubMed PMID: 32588343; PMCID: PMC8050124.

23. Miokovic T, Armbrecht G, Gast U, Rawer R, Roth HJ, Runge M, Felsenberg D, Belavy DL. Muscle atrophy, pain, and damage in bed rest reduced by resistive (vibration) exercise. Med Sci Sports Exerc. 2014;46(8):1506–16. doi: 10.1249/MSS.0000000000000279. PubMed PMID: 24561811.

24. Sicari BM, Agrawal V, Siu BF, Medberry CJ, Dearth CL, Turner NJ, Badylak SF. A murine model of volumetric muscle loss and a regenerative medicine approach for tissue replacement. Tissue Eng Part A. 2012;18(19-20):1941–8. doi: 10.1089/ten.TEA.2012.0475. PubMed PMID: 22906411; PMCID: PMC3463275.

25. Tejedera-Villafranca A, Montolio M, Ramon-Azcon J, Fernandez-Costa JM. Mimicking sarcolemmal damagein vitro: a contractile 3D model of skeletal muscle for drug testing in Duchenne muscular dystrophy. Biofabrication. 2023;15(4). Epub 20230927. doi: 10.1088/1758-5090/acfb3d. PubMed PMID: 37725998.

26. Jacho D, Yildirim-Ayan E. Mechanome-Guided Strategies in Regenerative Rehabilitation. Curr Opin Biomed Eng. 2024;29. Epub 20231130. doi: 10.1016/j.cobme.2023.100516. PubMed PMID: 38586151; PMCID: PMC10993906.

27. Kim JT, Kasukonis B, Dunlap G, Perry R, Washington T, Wolchok JC. Regenerative Repair of Volumetric Muscle Loss Injury is Sensitive to Age. Tissue Eng Part A. 2020;26(1-2):3–14. Epub 20190809. doi: 10.1089/ten.TEA.2019.0034. PubMed PMID: 31064280; PMCID: PMC6983754.

28. Hackmann MJ, Elliot JG, Green FHY, Cairncross A, Cense B, McLaughlin RA, Langton D, James AL, Noble PB, Donovan GM. Requirements and limitations of imaging airway smooth muscle throughout the lung in vivo. Respir Physiol Neurobiol. 2022;301:103884. Epub 20220315. doi: 10.1016/j.resp.2022.103884. PubMed PMID: 35301143.

29. Werkhausen A, Gloersen O, Nordez A, Paulsen G, Bojsen-Moller J, Seynnes OR. Linking muscle architecture and function in vivo: conceptual or methodological limitations? PeerJ. 2023;11:e15194. Epub 20230414. doi: 10.7717/peerj.15194. PubMed PMID: 37077309; PMCID: PMC10108853.

30. Timson BF. Evaluation of animal models for the study of exercise-induced muscle enlargement. J Appl Physiol (1985). 1990;69(6):1935–45. doi: 10.1152/jappl.1990.69.6.1935. PubMed PMID: 2076987.

31. Partridge T. Animal models of muscular dystrophy--what can they teach us? Neuropathol Appl Neurobiol. 1991;17(5):353–63. doi: 10.1111/j.1365-2990.1991.tb00735.x. PubMed PMID: 1758568.

32. Bernabei M, Lee SSM, Perreault EJ, Sandercock TG. Shear wave velocity is sensitive to changes in muscle stiffness that occur independently from changes in force. J Appl Physiol (1985). 2020;128(1):8–16. Epub 20190926. doi: 10.1152/japplphysiol.00112.2019. PubMed PMID: 31556833; PMCID: PMC6985815.

33. Smith LR, Meyer GA. Skeletal muscle explants: ex-vivo models to study cellular behavior in a complex tissue environment. Connect Tissue Res. 2020;61(3-4):248-61. Epub 2019/09/08. doi: 10.1080/03008207.2019.1662409. PubMed PMID: 31492079; PMCID: PMC8837600.

34. Naik NN, Vadloori B, Poosala S, Srivastava P, Coecke S, Smith A, Akhtar A, Roper C, Radhakrishnan S, Bhyravbhatla B, Damle M, Pulla VK, Hackethal J, Horland R, Li AP, Pati F, Singh MS, Occhetta P, Bisht R, Dandekar P, Bhagavatula K, Pajkrt D, Johnson M, Weber T, Huang J, Hysenaj L, Mallar B, Ramray B, Dixit S, Joshi S, Kulkarni M. Advances in Animal Models and Cutting-Edge Research in Alternatives: Proceedings of the Third International Conference on 3Rs Research and Progress, Vishakhapatnam, 2022. Altern Lab Anim. 2023;51(4):263–88. Epub 2023/06/07. doi: 10.1177/02611929231180428. PubMed PMID: 37282515.

35. MacArthur Clark J. The 3Rs in research: a contemporary approach to replacement, reduction and refinement. Br J Nutr. 2018;120(1):S1-S7. Epub 2017/10/31. doi: 10.1017/S0007114517002227. PubMed PMID: 29081302.

36. Szczesny SE. Ex vivo models of musculoskeletal tissues. Connect Tissue Res. 2020;61(3-4):245-7. Epub 2020/04/29. doi: 10.1080/03008207.2020.1742418. PubMed PMID: 32340565.

37. Caetano-Silva SP, Novicky A, Javaheri B, Rawlinson SCF, Pitsillides AA. Using Cell and Organ Culture Models to Analyze Responses of Bone Cells to Mechanical Stimulation. Methods Mol Biol. 2019;1914:99 – 128. doi: 10.1007/978-1-4939-8997-3_6. PubMed PMID: 30729462.

38. Ingersoll T, Cole S, Madren-Whalley J, Booker L, Dorsey R, Li A, Salem H. Generalized Additive Mixed-Models for Pharmacology Using Integrated Discrete Multiple Organ Co-Culture. PLoS One. 2016;11(4):e0152985. Epub 20160425. doi: 10.1371/journal.pone.0152985. PubMed PMID: 27110941; PMCID: PMC4844122.

39. Secerovic A, Ristaniemi A, Cui S, Li Z, Soubrier A, Alini M, Ferguson SJ, Weder G, Heub S, Ledroit D, Grad S. Toward the Next Generation of Spine Bioreactors: Validation of an Ex Vivo Intervertebral Disc Organ Model and Customized Specimen Holder for Multiaxial Loading. ACS Biomater Sci Eng. 2022;8(9):3969–76. Epub 20220817. doi: 10.1021/acsbiomaterials.2c00330. PubMed PMID: 35977717; PMCID: PMC9472220.

40. Christiansen BA, Bayly PV, Silva MJ. Constrained tibial vibration in mice: a method for studying the effects of vibrational loading of bone. J Biomech Eng. 2008;130(4):044502. doi: 10.1115/1.2917435. PubMed PMID: 18601464; PMCID: PMC2893880.

41. Smith EL, Kanczler JM, Oreffo RO. A new take on an old story: chick limb organ culture for skeletal niche development and regenerative medicine evaluation. Eur Cell Mater. 2013;26:91-106; discussion Epub 20130911. doi: 10.22203/ecm.v026a07. PubMed PMID: 24027022.

42. Viceconti M, Taddei F, Van Sint Jan S, Leardini A, Cristofolini L, Stea S, Baruffaldi F, Baleani M. Multiscale modelling of the skeleton for the prediction of the risk of fracture. Clin Biomech (Bristol, Avon). 2008;23(7):845–52. Epub 20080304. doi: 10.1016/j.clinbiomech.2008.01.009. PubMed PMID: 18304710.

43. Sargent M, Wark AW, Day S, Buis A. An ex vivo animal model to study the effect of transverse mechanical loading on skeletal muscle. Commun Biol. 2024;7(1):302. Epub 20240309. doi: 10.1038/s42003-024-05994-0. PubMed PMID: 38461200; PMCID: PMC10925026.

44. Carriero A, Abela L, Pitsillides AA, Shefelbine SJ. Ex vivo determination of bone tissue strains for an in vivo mouse tibial loading model. J Biomech. 2014;47(10):2490–7. Epub 20140403. doi: 10.1016/j.jbiomech.2014.03.035. PubMed PMID: 24835472; PMCID: PMC4071445.

45. Saito T, Nakamichi R, Yoshida A, Hiranaka T, Okazaki Y, Nezu S, Matsuhashi M, Shimamura Y, Furumatsu T, Nishida K, Ozaki T. The effect of mechanical stress on enthesis homeostasis in a rat Achilles enthesis organ culture model. J Orthop Res. 2021. Epub 20211115. doi: 10.1002/jor.25210. PubMed PMID: 34783068.

46. Geraldes DM, Modenese L, Phillips AT. Consideration of multiple load cases is critical in modelling orthotropic bone adaptation in the femur. Biomech Model Mechanobiol. 2016;15(5):1029–42. Epub 20151117. doi: 10.1007/s10237-015-0740-7. PubMed PMID: 26578078; PMCID: PMC5021760.

47. Shi M, Ishikawa M, Kamei N, Nakasa T, Adachi N, Deie M, Asahara T, Ochi M. Acceleration of skeletal muscle regeneration in a rat skeletal muscle injury model by local injection of human peripheral blood-derived CD133-positive cells. Stem Cells. 2009;27(4):949–60. doi: 10.1002/stem.4. PubMed PMID: 19353523.

48. Alheib O, da Silva LP, da Silva Morais A, Mesquita KA, Pirraco RP, Reis RL, Correlo VM. Injectable laminin-biofunctionalized gellan gum hydrogels loaded with myoblasts for skeletal muscle regeneration. Acta Biomater. 2022. Epub 20220309. doi: 10.1016/j.actbio.2022.03.008. PubMed PMID: 35278687.

49. Schaefer B, Beier JP, Ruhl T. Mesenchymal Stem Cells and the Generation of Neomuscle Tissue. Surg Technol Int. 2020;36:41-7. PubMed PMID: 32243565.

50. Miller BF, Hamilton KL, Majeed ZR, Abshire SM, Confides AL, Hayek AM, Hunt ER, Shipman P, Peelor FF, 3rd, Butterfield TA, Dupont-Versteegden EE. Enhanced skeletal muscle regrowth and remodelling in massaged and contralateral non-massaged hindlimb. J Physiol. 2018;596(1):83–103. Epub 20171201. doi: 10.1113/JP275089. PubMed PMID: 29090454; PMCID: PMC5746529.

51. Cenni V, Evangelisti C, Santi S, Sabatelli P, Neri S, Cavallo M, Lattanzi G, Mattioli E. Desmin and Plectin Recruitment to the Nucleus and Nuclei Orientation Are Lost in Emery-Dreifuss Muscular Dystrophy Myoblasts Subjected to Mechanical Stimulation. Cells. 2024;13(2). Epub 20240116. doi: 10.3390/cells13020162. PubMed PMID: 38247853; PMCID: PMC10814836.

52. Hakibilen C, Delort F, Daher MT, Joanne P, Cabet E, Cardoso O, Bourgois-Rocha F, Tian C, Rivas E, Madruga M, Ferreiro A, Lilienbaum A, Vicart P, Agbulut O, Henon S, Batonnet-Pichon S. Desmin Modulates Muscle Cell Adhesion and Migration. Front Cell Dev Biol. 2022;10:783724. Epub 20220308. doi: 10.3389/fcell.2022.783724. PubMed PMID: 35350386; PMCID: PMC8957967.

53. Krawetz RJ, Abubacker S, Leonard C, Masson AO, Shah S, Narendran N, Tailor P, Regmi SC, Labit E, Ninkovic N, Corpuz JM, Ito K, Underhill TM, Salo PT, Schmidt TA, Biernaskie JA. Proteoglycan 4 (PRG4) treatment enhances wound closure and tissue regeneration. NPJ Regen Med. 2022;7(1):32. Epub 20220624. doi: 10.1038/s41536-022-00228-5. PubMed PMID: 35750773; PMCID: PMC9232611.

54. De Croos JN, Dhaliwal SS, Grynpas MD, Pilliar RM, Kandel RA. Cyclic compressive mechanical stimulation induces sequential catabolic and anabolic gene changes in chondrocytes resulting in increased extracellular matrix accumulation. Matrix Biol. 2006;25(6):323–31. Epub 2006/05/16. doi: 10.1016/j.matbio.2006.03.005. PubMed PMID: 16697175.

55. Jin M, Frank EH, Quinn TM, Hunziker EB, Grodzinsky AJ. Tissue shear deformation stimulates proteoglycan and protein biosynthesis in bovine cartilage explants. Arch Biochem Biophys. 2001;395(1):41 – 8. doi: 10.1006/abbi.2001.2543. PubMed PMID: 11673864.

56. Costamagna D, Duelen R, Penna F, Neumann D, Costelli P, Sampaolesi M. Interleukin-4 administration improves muscle function, adult myogenesis, and lifespan of colon carcinoma-bearing mice. J Cachexia Sarcopenia Muscle. 2020;11(3):783–801. Epub 20200227. doi: 10.1002/jcsm.12539. PubMed PMID: 32103619; PMCID: PMC7296260.

57. Jacho D, Rabino A, Garcia-Mata R, Yildirim-Ayan E. Mechanoresponsive regulation of fibroblast-to-myofibroblast transition in three-dimensional tissue analogues: mechanical strain amplitude dependency of fibrosis. Sci Rep. 2022;12(1):16832. Epub 20221007. doi: 10.1038/s41598-022-20383-5. PubMed PMID: 36207437; PMCID: PMC9547073.

58. Chen J, Zhou R, Feng Y, Cheng L. Molecular mechanisms of exercise contributing to tissue regeneration. Signal Transduct Target Ther. 2022;7(1):383. Epub 20221130. doi: 10.1038/s41392-022-01233-2. PubMed PMID: 36446784; PMCID: PMC9709153.

59. Discher DE, Janmey P, Wang YL. Tissue cells feel and respond to the stiffness of their substrate. Science. 2005;310(5751):1139–43. doi: 10.1126/science.1116995. PubMed PMID: 16293750.

60. Kirk B, Duque G. Muscle and Bone: An Indissoluble Union. J Bone Miner Res. 2022;37(7):1211–2. Epub 20220628. doi: 10.1002/jbmr.4626. PubMed PMID: 35764095.

61. Tanaka Y, Kita S, Nishizawa H, Fukuda S, Fujishima Y, Obata Y, Nagao H, Masuda S, Nakamura Y, Shimizu Y, Mineo R, Natsukawa T, Funahashi T, Ranscht B, Fukada SI, Maeda N, Shimomura I. Author Correction: Adiponectin promotes muscle regeneration through binding to T-cadherin. Sci Rep. 2020;10(1):12219. Epub 20200717. doi: 10.1038/s41598-020-66545-1. PubMed PMID: 32678119; PMCID: PMC7367247.

62. De Filippo K, Dudeck A, Hasenberg M, Nye E, van Rooijen N, Hartmann K, Gunzer M, Roers A, Hogg N. Mast cell and macrophage chemokines CXCL1/CXCL2 control the early stage of neutrophil recruitment during tissue inflammation. Blood. 2013;121(24):4930–7. Epub 20130503. doi: 10.1182/blood-2013-02-486217. PubMed PMID: 23645836.

63. Zhang L, Ran L, Garcia GE, Wang XH, Han S, Du J, Mitch WE. Chemokine CXCL16 regulates neutrophil and macrophage infiltration into injured muscle, promoting muscle regeneration. Am J Pathol. 2009;175(6):2518–27. Epub 20091105. doi: 10.2353/ajpath.2009.090275. PubMed PMID: 19893053; PMCID: PMC2789607.

64. Conover CA. Insulin-like growth factor-binding proteins and bone metabolism. Am J Physiol Endocrinol Metab. 2008;294(1):E10–4. Epub 20071114. doi: 10.1152/ajpendo.00648.2007. PubMed PMID: 18003717.

65. Shinde AV, Humeres C, Frangogiannis NG. The role of alpha-smooth muscle actin in fibroblast-mediated matrix contraction and remodeling. Biochim Biophys Acta Mol Basis Dis. 2017;1863(1):298 –309. Epub 20161104. doi: 10.1016/j.bbadis.2016.11.006. PubMed PMID: 27825850; PMCID: PMC5163362.

66. Shintaku J, Peterson JM, Talbert EE, Gu JM, Ladner KJ, Williams DR, Mousavi K, Wang R, Sartorelli V, Guttridge DC. MyoD Regulates Skeletal Muscle Oxidative Metabolism Cooperatively with Alternative NF-kappaB. Cell Rep. 2016;17(2):514–26. doi: 10.1016/j.celrep.2016.09.010. PubMed PMID: 27705798; PMCID: PMC5059110.

67. von Maltzahn J, Jones AE, Parks RJ, Rudnicki MA. Pax7 is critical for the normal function of satellite cells in adult skeletal muscle. Proc Natl Acad Sci U S A. 2013;110(41):16474–9. Epub 20130924. doi: 10.1073/pnas.1307680110. PubMed PMID: 24065826; PMCID: PMC3799311.

68. Moretti I, Ciciliot S, Dyar KA, Abraham R, Murgia M, Agatea L, Akimoto T, Bicciato S, Forcato M, Pierre P, Uhlenhaut NH, Rigby PW, Carvajal JJ, Blaauw B, Calabria E, Schiaffino S. MRF4 negatively regulates adult skeletal muscle growth by repressing MEF2 activity. Nat Commun. 2016;7:12397. Epub 20160803. doi: 10.1038/ncomms12397. PubMed PMID: 27484840; PMCID: PMC4976255.

69. de Araujo Farias V, Carrillo-Galvez AB, Martin F, Anderson P. TGF-beta and mesenchymal stromal cells in regenerative medicine, autoimmunity and cancer. Cytokine Growth Factor Rev. 2018;43:25–37. Epub 20180613. doi: 10.1016/j.cytogfr.2018.06.002. PubMed PMID: 29954665.

70. Huang S, Thomsson KA, Jin C, Ryberg H, Das N, Struglics A, Rolfson O, Bjorkman LI, Eisler T, Schmidt TA, Jay GD, Krawetz R, Karlsson NG. Truncated lubricin glycans in osteoarthritis stimulate the synoviocyte secretion of VEGFA, IL-8, and MIP-1alpha: Interplay between O-linked glycosylation and inflammatory cytokines. Front Mol Biosci. 2022;9:942406. Epub 20220921. doi: 10.3389/fmolb.2022.942406. PubMed PMID: 36213120; PMCID: PMC9532613.

71. Bock F, Maruyama K, Regenfuss B, Hos D, Steven P, Heindl LM, Cursiefen C. Novel anti(lymph)angiogenic treatment strategies for corneal and ocular surface diseases. Prog Retin Eye Res. 2013;34:89–124. Epub 20130122. doi: 10.1016/j.preteyeres.2013.01.001. PubMed PMID: 23348581.

72. Jang DI, Lee AH, Shin HY, Song HR, Park JH, Kang TB, Lee SR, Yang SH. The Role of Tumor Necrosis Factor Alpha (TNF-alpha) in Autoimmune Disease and Current TNF-alpha Inhibitors in Therapeutics. Int J Mol Sci. 2021;22(5). Epub 20210308. doi: 10.3390/ijms22052719. PubMed PMID: 33800290; PMCID: PMC7962638.

73. Lopez-Castejon G, Brough D. Understanding the mechanism of IL-1beta secretion. Cytokine Growth Factor Rev. 2011;22(4):189–95. Epub 20111022. doi: 10.1016/j.cytogfr.2011.10.001. PubMed PMID: 22019906; PMCID: PMC3714593.

74. Etzerodt A, Moestrup SK. CD163 and inflammation: biological, diagnostic, and therapeutic aspects. Antioxid Redox Signal. 2013;18(17):2352–63. Epub 20121019. doi: 10.1089/ars.2012.4834. PubMed PMID: 22900885; PMCID: PMC3638564.

75. Casarosa P, Waldhoer M, LiWang PJ, Vischer HF, Kledal T, Timmerman H, Schwartz TW, Smit MJ, Leurs R. CC and CX3C chemokines differentially interact with the N terminus of the human cytomegalovirus-encoded US28 receptor. J Biol Chem. 2005;280(5):3275–85. Epub 20041116. doi: 10.1074/jbc.M407536200. PubMed PMID: 15546882.

76. Kiani AK, Pheby D, Henehan G, Brown R, Sieving P, Sykora P, Marks R, Falsini B, Capodicasa N, Miertus S, Lorusso L, Dondossola D, Tartaglia GM, Ergoren MC, Dundar M, Michelini S, Malacarne D, Bonetti G, Dautaj A, Donato K, Medori MC, Beccari T, Samaja M, Connelly ST, Martin D, Morresi A, Bacu A, Herbst KL, Kapustin M, Stuppia L, Lumer L, Farronato G, Bertelli M, International Bioethics Study G. Ethical considerations regarding animal experimentation. J Prev Med Hyg. 2022;63(2 Suppl 3):E255-E66. Epub 2022/12/09. doi: 10.15167/2421-4248/jpmh2022.63.2S3.2768. PubMed PMID: 36479489; PMCID: PMC9710398.

77. Kirk RGW. Recovering The Principles of Humane Experimental Technique: The 3Rs and the Human Essence of Animal Research. Sci Technol Human Values. 2018;43(4):622–48. Epub 2018/07/17. doi: 10.1177/0162243917726579. PubMed PMID: 30008492; PMCID: PMC6027778.

78. Diaz L, Zambrano E, Flores ME, Contreras M, Crispin JC, Aleman G, Bravo C, Armenta A, Valdes VJ, Tovar A, Gamba G, Barrios-Payan J, Bobadilla NA. Ethical Considerations in Animal Research: The Principle of 3R’s. Rev Invest Clin. 2020;73(4):199–209. Epub 2020/10/23. doi: 10.24875/RIC.20000380. PubMed PMID: 33090120.

79. Azizi P, Drobek C, Budday S, Seitz H. Simulating the mechanical stimulation of cells on a porous hydrogel scaffold using an FSI model to predict cell differentiation. Front Bioeng Biotechnol. 2023;11:1249867. Epub 20230919. doi: 10.3389/fbioe.2023.1249867. PubMed PMID: 37799813; PMCID: PMC10549991.

80. Pedaprolu K, Szczesny S. A Novel, Open Source, Low-Cost Bioreactor for Load-Controlled Cyclic Loading of Tendon Explants. J Biomech Eng. 2022. Epub 20220211. doi: 10.1115/1.4053795. PubMed PMID: 35147179.

81. Subramanian G, Elsaadany M, Bialorucki C, Yildirim-Ayan E. Creating homogenous strain distribution within 3D cell-encapsulated constructs using a simple and cost-effective uniaxial tensile bioreactor: Design and validation study. Biotechnol Bioeng. 2017;114(8):1878–87. Epub 2017/04/21. doi: 10.1002/bit.26304. PubMed PMID: 28425561.

82. Warden SJ. Animal models for the study of tendinopathy. Br J Sports Med. 2007;41(4):232–40. Epub 20061124. doi: 10.1136/bjsm.2006.032342. PubMed PMID: 17127722; PMCID: PMC2658951.

83. Palmes D, Spiegel HU, Schneider TO, Langer M, Stratmann U, Budny T, Probst A. Achilles tendon healing: long-term biomechanical effects of postoperative mobilization and immobilization in a new mouse model. J Orthop Res. 2002;20(5):939–46. doi: 10.1016/S0736-0266(02)00032-3. PubMed PMID: 12382957.

84. Yu Y, Li K, Peng Y, Wu W, Chen F, Shao Z, Zhang Z. Animal models of cancer metastasis to the bone. Front Oncol. 2023;13:1165380. Epub 20230405. doi: 10.3389/fonc.2023.1165380. PubMed PMID: 37091152; PMCID: PMC10113496.

85. Baroi S, Czernik PJ, Chougule A, Griffin PR, Lecka-Czernik B. PPARG in osteocytes controls sclerostin expression, bone mass, marrow adiposity and mediates TZD-induced bone loss. Bone. 2021;147:115913. Epub 20210316. doi: 10.1016/j.bone.2021.115913. PubMed PMID: 33722775; PMCID: PMC8076091.

86. Bouxsein ML, Boyd SK, Christiansen BA, Guldberg RE, Jepsen KJ, Muller R. Guidelines for assessment of bone microstructure in rodents using micro-computed tomography. J Bone Miner Res. 2010;25(7):1468-doi: 10.1002/jbmr.141. PubMed PMID: 20533309.

87. Baylan N, Bhat S, Ditto M, Lawrence JG, Lecka-Czernik B, Yildirim-Ayan E. Polycaprolactone nanofiber interspersed collagen type-I scaffold for bone regeneration: a unique injectable osteogenic scaffold. Biomed Mater. 2013;8(4):045011. Epub 2013/06/28. doi: 10.1088/1748-6041/8/4/045011. PubMed PMID: 23804651.

88. Subramanian G, Bialorucki C, Yildirim-Ayan E. Nanofibrous yet injectable polycaprolactone-collagen bone tissue scaffold with osteoprogenitor cells and controlled release of bone morphogenetic protein-2. Mater Sci Eng C Mater Biol Appl. 2015;51:16–27. Epub 2015/04/07. doi: 10.1016/j.msec.2015.02.030. PubMed PMID: 25842103.

89. Shortridge C, Akbari Fakhrabadi E, Wuescher LM, Worth RG, Liberatore MW, Yildirim-Ayan E. Impact of Digestive Inflammatory Environment and Genipin Crosslinking on Immunomodulatory Capacity of Injectable Musculoskeletal Tissue Scaffold. Int J Mol Sci. 2021;22(3). Epub 2021/01/28. doi: 10.3390/ijms22031134. PubMed PMID: 33498864; PMCID: PMC7866115.

90. Roebke E, Jacho D, Eby O, Aldoohan S, Elsamaloty H, Yildirim-Ayan E. Injectable Cell-Laden Nanofibrous Matrix for Treating Annulus Fibrosus Defects in Porcine Model: An Organ Culture Study. Life (Basel). 2022;12(11). Epub 2022/11/27. doi: 10.3390/life12111866. PubMed PMID: 36431001; PMCID: PMC9694927.

